# DRP1 Orchestrates RNA Virus Permissiveness Through Metabolic Reprogramming Independent of Mitochondrial Dynamics

**DOI:** 10.64898/2026.11.19.767996

**Authors:** Poojitha Sai Potharaju, Jayasurya Doddetipalli, Vishal Sah, Prangya Paramita Sahoo, Aswathi Chandran, Dixit Tandel, Malabika Bhowmik, Jhansi Vadlamudi, Priyanka Pandey, Krishnan H Harshan

## Abstract

Viruses reprogram host metabolism and mitochondrial dynamics to support replication, yet how these processes govern viral permissiveness remains unclear. Using SARS-CoV-2 alongside other RNA viruses JEV, VSV, and HCV, we identify DRP1 as a central determinant of the virus-permissive cellular state beyond mitochondrial fission. While RNA viruses induce pronounced mitochondrial fragmentation linked to enhanced permissiveness, MFN1 conferred potent antiviral protection independent of type I interferon. Targeting DRP1-associated functions pharmacologically suppresses viral replication, accompanied by decreased glycolysis and enhanced oxidative metabolism. Remarkably, genetic depletion of DRP1 itself potently restricts viral replication in both wild-type and MFN1-deficient cells despite persistent mitochondrial fragmentation in the latter, demonstrating that MFN1 and mitochondrial elongation are dispensable for antiviral restriction. Together, these findings identify DRP1 as a key regulator of viral permissiveness operating beyond mitochondrial morphology. By uncoupling mitochondrial morphology from antiviral regulation, our study highlights DRP1-associated pathways as promising targets for host-directed antiviral strategies.

**HIGHLIGHTS:** RNA viruses activate DRP1 and promote mitochondrial fragmentation

Pharmacological and genetic targeting of DRP1 function potently restricts viral replication

MFN1 and mitochondrial fusion are dispensable for DRP1-associated viral restriction

DRP1 regulates RNA virus permissiveness through metabolic reprogramming

## INTRODUCTION

Viruses are obligate intracellular pathogens that depend extensively on host-cell machinery for genome replication, protein synthesis, and progeny production. Successful infection therefore requires remodeling of cellular pathways involved in innate immune signaling, organelle homeostasis, and metabolism^1^. During RNA virus infection, cytosolic viral RNA is primarily detected by RIG-I-like receptors RIG-I and MDA5, which signal through the mitochondrial antiviral-signaling protein (MAVS) to activate type I interferon (IFN) and pro-inflammatory responses^2,3^. The identification of MAVS established mitochondria as important platforms for antiviral signaling and highlighted a broader connection between mitochondrial homeostasis and innate immunity^2,4,5^. Beyond their role in oxidative metabolism and ATP production, mitochondria regulate apoptosis, calcium and redox homeostasis, metabolic adaptation, and immune responses^4^. Several components of innate immune signaling are functionally associated with mitochondria, allowing changes in mitochondrial homeostasis during infection to influence both immune signaling and the metabolic environment that supports viral replication^2,5,6^.

A central determinant of mitochondrial homeostasis is the balance between mitochondrial fusion and fission. Fusion is mediated by GTPases mitofusin 1 (MFN1) and mitofusin 2 (MFN2) at the outer mitochondrial membrane and another GTPase OPA1 at the inner mitochondrial membrane, promoting mitochondrial content mixing and network connectivity. In contrast, mitochondrial fission is primarily controlled by the dynamin-related GTPase DRP1, which is recruited to the outer mitochondrial membrane through receptors including Mitochondrial Fission Factor (MFF), FIS1, MiD49, and MiD51, where it oligomerizes to constrict and divide the organelle^7,8^. DRP1 activity is dynamically regulated by post-translational modifications, particularly phosphorylation. Phosphorylation at Ser616 can be mediated by several kinases, including CDK1/5, ERK1/2, PKCδ, and CaMKII, thereby regulating DRP1 activity in a stimulus-dependent manner^9–13^. Mitochondrial dynamics are also closely integrated with quality control: fission facilitates segregation and mitophagic removal of dysfunctional mitochondrial regions, whereas fusion promotes functional complementation and preservation of mitochondrial integrity during stress. Thus, fusion, fission, and mitophagy constitute an interconnected network regulating mitochondrial architecture and function to changing cellular demands. Perturbation of this balance can have consequences extending beyond mitochondrial morphology, including alterations in mitochondrial signaling and quality control^7,14^.

Mitochondrial dynamics are also tightly connected to cellular metabolism. Changes in mitochondrial architecture can accompany transitions between glycolytic and oxidative metabolic states, although these relationships are highly dependent on cellular context. Increased mitochondrial fragmentation is frequently associated with enhanced glycolytic metabolism, whereas interconnected mitochondrial networks can support mitochondrial respiration and oxidative metabolism^15,16^. Such metabolic plasticity is particularly relevant during viral infection. Rapidly replicating viruses can exploit enhanced glycolytic flux to obtain ATP and biosynthetic intermediates required for nucleotide, lipid, and amino-acid synthesis^1,16,17^. Conversely, metabolic states that limit glycolytic availability or preserve mitochondrial oxidative metabolism can restrict replication in several viral systems^16–18^. Mitochondrial dynamics may therefore provide an important interface between organelle architecture and the metabolic state of infected cells^15,16^.

Viruses frequently manipulate mitochondrial dynamics, although the direction and functional consequences of this remodeling vary among viral systems^16,19–23^. Hepatitis C virus, for example, promotes DRP1-dependent mitochondrial fragmentation and PARKIN-associated mitophagy^19^, whereas dengue virus has been reported in specific experimental settings to suppress DRP1 recruitment and promote mitochondrial elongation^20,21^. Such divergent strategies suggest that mitochondrial remodeling is not merely a consequence of infection-associated stress but may contribute to establishing cellular conditions favorable for viral replication, immune evasion, or metabolic adaptation ^16,19–23^.

Mitochondrial dynamics can also influence MAVS-dependent IFN signaling, with mitochondrial elongation associated with enhanced antiviral signaling and fragmentation linked to impaired organization of mitochondrial signaling platforms^24^. However, whether mitochondrial remodeling itself determines cellular permissiveness to RNA-virus infection, and whether it can establish an antiviral state independently of canonical type I interferon signaling, remains incompletely understood. SARS-CoV-2, the causative agent of COVID-19, provides a relevant model to investigate these relationships, given its well-documented suppression of type I IFN activation^25,26^. Additionally, several SARS-CoV-2 proteins have been reported to interact with mitochondrial proteins or regulators of mitochondrial antiviral signaling ^27–30^, suggesting that manipulation of mitochondrial function is an important component of infection. Nevertheless, the functional relationships among SARS-CoV-2-induced mitochondrial remodeling, DRP1 activation, mitochondrial quality control, metabolic reprogramming, and antiviral signaling remain poorly defined.

Here, we investigated how mitochondrial dynamics and associated proteins influence cellular permissiveness to RNA-virus infection, using SARS-CoV-2 as the primary mechanistic model and extending key findings to JEV, HCV, and VSV. We observed that SARS-CoV-2 infection promoted DRP1- associated mitochondrial fission, followed by mitochondrial fragmentation and enhanced mitophagic signaling, with similar mitochondrial remodeling observed during infection with other RNA viruses. The functional significance of these changes became evident through modulation of mitochondrial dynamics-associated proteins, where MFN1 deficiency markedly enhanced viral replication, whereas MFN1 overexpression and interventions targeting DRP1-associated functions established potent antiviral protection independently of type I interferon signaling. Further investigation revealed a close association between these regulatory proteins and cellular metabolism, as MFN1 deficiency favored a glycolytic, virus-permissive environment, while interventions suppressing glycolysis or promoting mitochondrial pyruvate import were associated with reduced viral replication. Importantly, the antiviral effects of DRP1-targeted interventions, glycolytic suppression, and pyruvate supplementation persisted in MFN1-deficient cells despite sustained mitochondrial fragmentation, demonstrating that MFN1-mediated mitochondrial elongation is dispensable for establishing these antiviral effects. This distinction was further strengthened by genetic DRP1 depletion, which potently restricted viral replication even in the absence of MFN1. Moreover, the greater antiviral effects observed following DRP1 targeting compared with glycolytic inhibition alone suggest that DRP1 may influence viral permissiveness through additional cellular functions beyond glycolytic regulation. Together, our findings identify DRP1 as a central determinant of RNA-virus permissiveness, revealing a functional relationship between mitochondrial dynamics-associated proteins and metabolic regulation that extends beyond mitochondrial morphology and potentially provides new opportunities for host-directed antiviral intervention.

## RESULTS

Mitochondrial fission is an outcome of SARS-CoV-2 infection and a recurring feature of predominant RNA viral infections We investigated how SARS-CoV-2 infection influences the molecular processes that regulate mitochondrial dynamics. Previous studies, including our own, have demonstrated that SARS-CoV-2 promotes mitochondrial fragmentation and mitophagy^6,31–33^ (Figure S1A). To determine whether this morphological alteration is accompanied by activation of the mitochondrial fission machinery, we examined the temporal regulation of key proteins involved in mitochondrial dynamics. Phosphorylation of DRP1 at S616, an important marker of its fission-associated activity, showed a moderate increase as early as 6 hours post-infection (hpi), which became more pronounced by 24 hpi. Interestingly, DRP1 phosphorylation subsequently declined at 48 hpi and beyond, suggesting that fission-associated signaling is most prominent during the early stages of infection (Figures 1 A & B). Consistent with this observation, mitochondrial fission factor (MFF) levels increased during the first 24 hours of infection but declined markedly at later time points (Figures 1 A & C). In contrast, the mitochondrial fusion proteins MFN1 and MFN2 remained relatively unchanged during the initial 24 hours of infection but underwent substantial depletion thereafter (Figures 1 A, D, & E). Together, these observations suggest that SARS- CoV-2 infection promotes early activation of DRP1-associated mitochondrial fission, followed by a decline in the abundance of mitochondrial dynamics-associated proteins. To further verify these findings, we examined the mitochondrial association of phosphorylated DRP1 in SARS-CoV-2-infected Huh-7 cells using confocal microscopy. Mitochondria were visualized using MitoTracker Red, while phosphorylated DRP1 was detected by immunostaining. Consistent with the biochemical observations, SARS-CoV-2 infection significantly enhanced the mitochondrial association of phosphorylated DRP1, indicating increased recruitment of activated DRP1 to mitochondria. Interestingly, this association remained elevated during infection despite a substantial reduction in mitochondrial signals from 48 hpi onward (Figures 1 F-H). These results demonstrate that SARS-CoV-2 infection triggers persistent mitochondrial fission.

**Figure 1.**
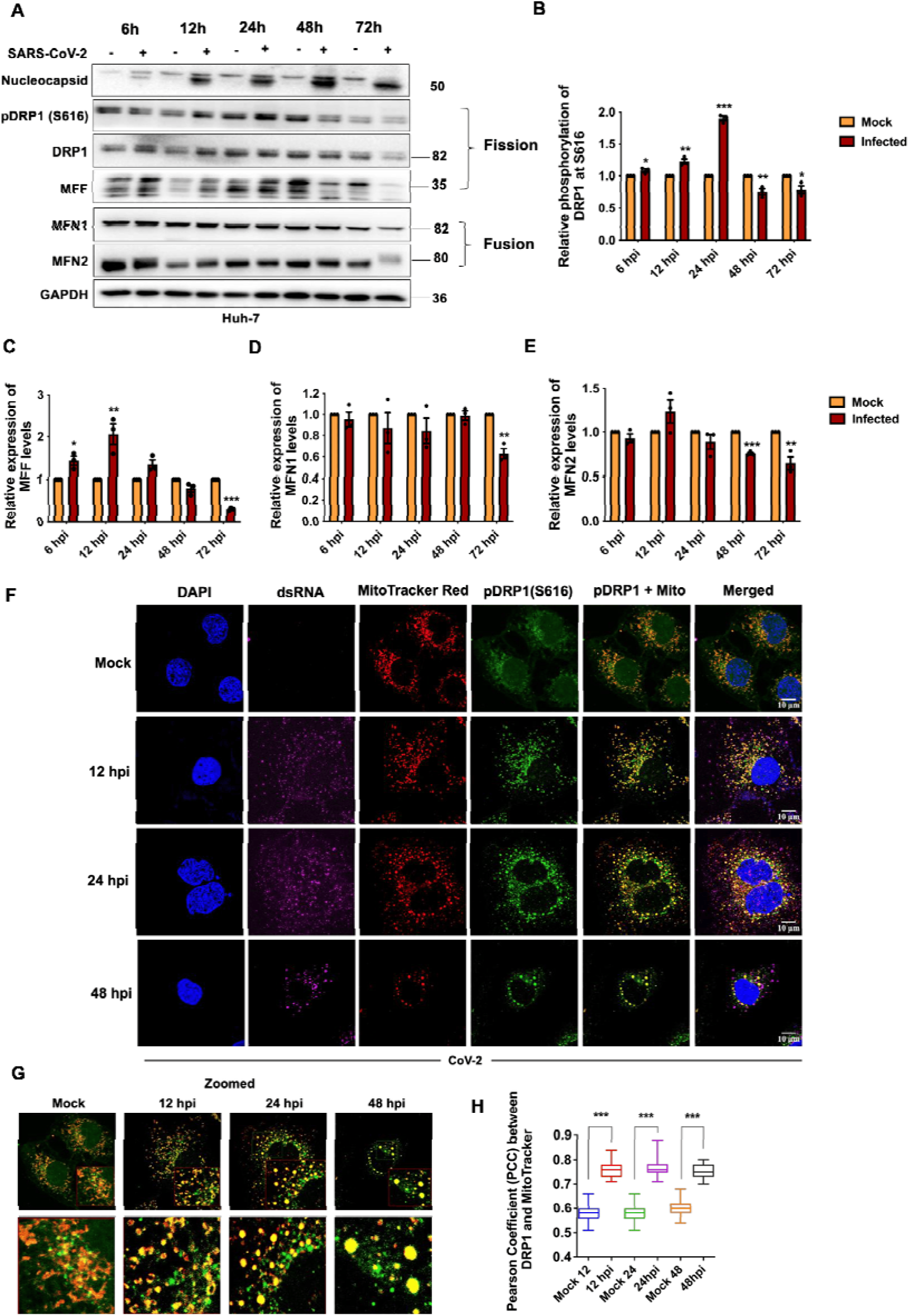
Mitochondrial fission is induced during SARS-CoV-2 infection and represents a recurring feature of RNA-virus infections. (A) Immunoblots showing SARS-CoV-2 Nucleocapsid protein and mitochondrial dynamics-associated proteins, including phosphorylated DRP1 (pDRP1-S616), total DRP1, MFF, MFN1, and MFN2, in Huh-7 cells at the indicated time points following SARS-CoV-2 infection. (B–E) Densitometric quantification of the indicated mitochondrial dynamics-associated proteins shown in (A). (F) Confocal microscopy images showing the mitochondrial localization of pDRP1-S616 (green) in SARS- CoV-2 infected Huh-7 cells at the indicated time points. Mitochondria were labeled with MitoTracker Red (red). dsRNA, stained in purple, confirms viral replication. (G) Magnified images of the co-localized regions in (F) (H) Quantification of pDRP1-S616 colocalization with MitoTracker Red in (F) using Pearson’s correlation coefficient n=250 ROIs. Scale bars, 10 µm and n = 50 cells unless otherwise indicated.

In our previous study, we demonstrated that SARS-CoV-2 infection induces pronounced mitophagy accompanied by significant depletion in mitochondrial mass by 48 hpi^6^. The decline in the abundance of both fission-associated proteins (DRP1 and MFF) and fusion proteins (MFN1 and MFN2) at later time points in the present study is consistent with this progressive loss of mitochondrial mass. In agreement with this interpretation, we noticed substantially elevated PINK1 phosphorylation beyond 48 hpi (Figures S1 B & C), indicating sustained activation of mitophagy. Collectively, these findings suggest that SARS-CoV-2 infection promotes mitochondrial fission as early as 6 hpi, characterized by increased DRP1 phosphorylation and MFF abundance, followed by enhanced mitophagy and mitochondrial depletion during the later stages of infection.

Given that mitochondrial fragmentation and mitophagy have been reported during several RNA-virus infections^16,19–21^, we next investigated whether similar mitochondrial remodeling occurs in response to other viruses in our experimental system. Consistent with previous observations, infection with JEV and HCV (positive-sense single-stranded RNA viruses) and VSV (a negative-sense single-stranded RNA virus) resulted in pronounced mitochondrial fragmentation (Figures S1 D & E). Together, these findings indicate that mitochondrial fragmentation is a recurring feature of RNA-virus infection and suggest that remodeling of the mitochondrial network may represent a common aspect of virus-host interactions.

### Mitochondrial fission associated with MFN1 depletion enhances viral permissiveness

Since mitochondrial fission is commonly observed during RNA-virus infections, we investigated whether mitochondrial fission provides a replicative advantage to the virus. To address this, we depleted MFN1 in Huh-7 cells using specific siRNA, followed by SARS-CoV-2 infection. Efficient MFN1 knockdown (KD) (Figures 2 A & B) resulted in a marked increase in SARS-CoV-2 Nucleocapsid protein levels accompanied by an approximately 2-log increase in viral titers (Figure 2 A & C). In contrast, MFN2 depletion resulted in a comparatively modest increase in SARS-CoV-2 replication (Figures 2 D-F), suggesting that MFN1 plays a more prominent role in restricting viral replication. These observations indicated that disruption of mitochondrial fusion through MFN1 depletion establishes a cellular environment more permissive to SARS-CoV-2 replication. To further substantiate these findings, we generated MFN1-knockout (KO) Huh- 7 cells using CRISPR-Cas9 and characterized two independently isolated KO clones (C1 and C2) alongside the pooled KO population. Both independent clones and the pooled KO population consistently exhibited fragmented mitochondrial morphology (Figures 2 G & H; Figures S2 A & B). Consistent with the observed mitochondrial fragmentation, MFN1 KO cells also exhibited significantly increased DRP1 phosphorylation (Figures 2 I & J). Both MFN1 KO clones and pooled cultures supported enhanced SARS- CoV-2 replication, as evidenced by elevated viral N protein levels (Figures 2K; Figures S2 C & D) and significantly increased viral titers (Figure 2L; Figure S2E). To determine whether this phenotype was conserved across different cellular backgrounds, we examined SARS-CoV-2 replication in MFN1 KO mouse embryonic fibroblasts (MEFs). We previously reported robust SARS-CoV-2 replication in MEFs^6^. Consistent with the findings in Huh-7 cells, MFN1 deficiency significantly enhanced SARS-CoV-2 replication in MEFs as well (Figures S2 F & G), further supporting a proviral phenotype associated with loss of MFN1. Moreover, MFN1 KO cells also supported increased JEV replication (Figures 2 M & N), indicating that the enhanced viral permissiveness associated with MFN1 deficiency is not restricted to SARS-CoV-2.

**Figure 2.**
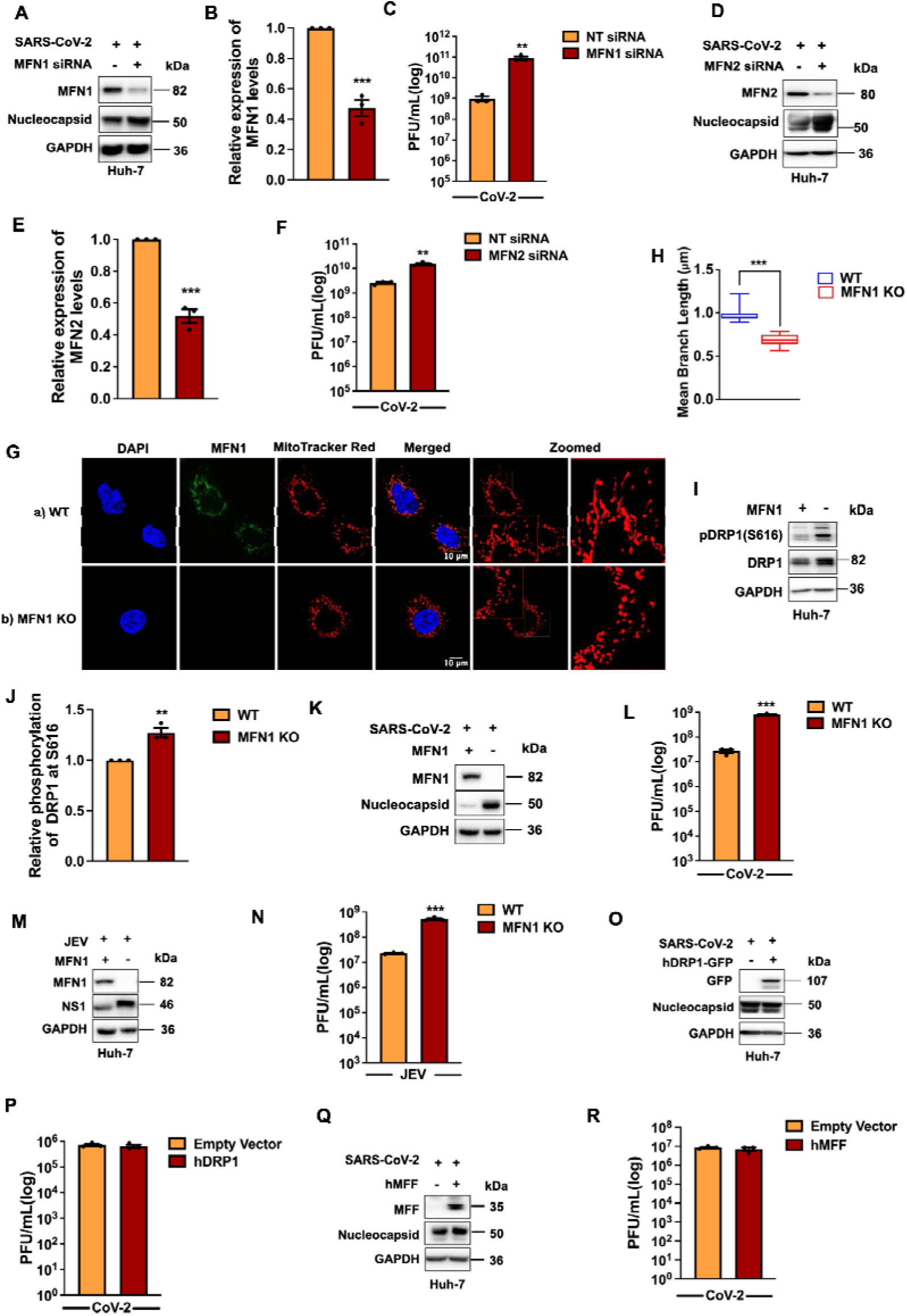
Mitochondrial fission associated with MFN1 depletion enhances viral permissiveness. (A) Immunoblots showing MFN1 and SARS-CoV-2 Nucleocapsid protein levels in Huh-7 cells transfected with MFN1-specific siRNA followed by SARS-CoV-2 infection. (B) Densitometric quantification of MFN1 protein levels shown in (A). (C) Infectious viral titers measured in culture supernatants from (A). (D) Immunoblots showing MFN2 and SARS-CoV-2 Nucleocapsid protein levels in Huh-7 cells transfected with MFN2-specific siRNA, followed by SARS-CoV-2 infection. (E) Densitometric quantification of MFN2 protein levels shown in (D). (F) Infectious viral titers measured in culture supernatants from (D). (G) Confocal microscopy images showing mitochondrial fragmentation in wild-type (WT) and MFN1- knockout (KO) Huh-7 cells. Clone C1 was used for the experiments shown in this panel and subsequent experiments unless otherwise indicated. Mitochondria were stained with MitoTracker Red (red) and MFN1 stained (green). (H) Quantification of mean mitochondrial branch length in cells shown in (G). (I) Immunoblots showing pDRP1-S616 in WT and MFN1 KO Huh-7 cells. (J) Densitometric quantification of relative DRP1 phosphorylation shown in (I). (K) Immunoblots showing MFN1 and SARS-CoV-2 Nucleocapsid protein levels in WT and MFN1 KO Huh-7 cells following SARS-CoV-2 infection for 24 h. (L) Infectious viral titers measured in culture supernatants from (K). (M) Immunoblots showing MFN1 and JEV NS1 protein levels in WT and MFN1 KO Huh-7 cells following JEV infection for 24 h. (N) Infectious viral titers measured in culture supernatants from (M). (O) Representative immunoblots showing GFP-tagged human DRP1 (hDRP1-GFP) expression and SARS- CoV-2 Nucleocapsid protein levels in Huh-7 cells transfected with hDRP1-GFP followed by SARS-CoV-2 infection. (P) Infectious viral titers measured in culture supernatants from (O). (Q) Immunoblots showing MFF and SARS-CoV-2 Nucleocapsid protein levels in Huh-7 cells transfected with human MFF followed by SARS-CoV-2 infection. (R) Infectious viral titers measured in culture supernatants from (Q). Scale bars, 10 µm. n = 50 cells unless otherwise indicated.

We next examined whether augmenting the mitochondrial fission machinery could further enhance viral replication. Interestingly, overexpression of neither DRP1-eGFP nor MFF could further increase SARS- CoV-2 replication (Figures 2 O-R), despite their established roles in mitochondrial fission. This observation raises the possibility that SARS-CoV-2 infection already engages the mitochondrial fission machinery to its fullest capacity, such that additional expression of its components does not further enhance viral replication. Collectively, these findings demonstrate that MFN1 deficiency establishes a cellular environment favorable for RNA-virus replication, supporting a strong association between impaired mitochondrial fusion, increased mitochondrial fragmentation, and enhanced viral permissiveness.

### Mitochondrial fusion establishes an antiviral state in host cells independent of type I interferon signaling

Since mitochondrial fission established a more robust environment for viral replication, we next investigated whether promoting mitochondrial fusion could establish an antiviral state. It is well established that inhibition of mitochondrial fission through DRP1 depletion promotes mitochondrial elongation^34,35^. Therefore, we initially examined the effect of DRP1 knockdown (KD) on SARS-CoV-2 replication. DRP1 depletion resulted in robust suppression of viral replication (Figures 3 A-C), supporting an antiviral effect associated with inhibition of mitochondrial fission. We subsequently investigated whether promoting mitochondrial fusion through MFN1 overexpression could similarly restrict viral replication. As expected, MFN1 overexpression resulted in pronounced mitochondrial elongation in Huh- 7 cells (Supplementary Figures 3 A & B). Upon SARS-CoV-2 infection, MFN1-overexpressing cells maintained an elongated mitochondrial network, in contrast to the extensive fragmentation observed in empty vector control cells (Figures 3 D & E). Importantly, MFN1 overexpression markedly suppressed SARS-CoV-2 replication, as evidenced by reduced N protein levels and viral titers (Figures 3 F & G).

**Figure 3.**
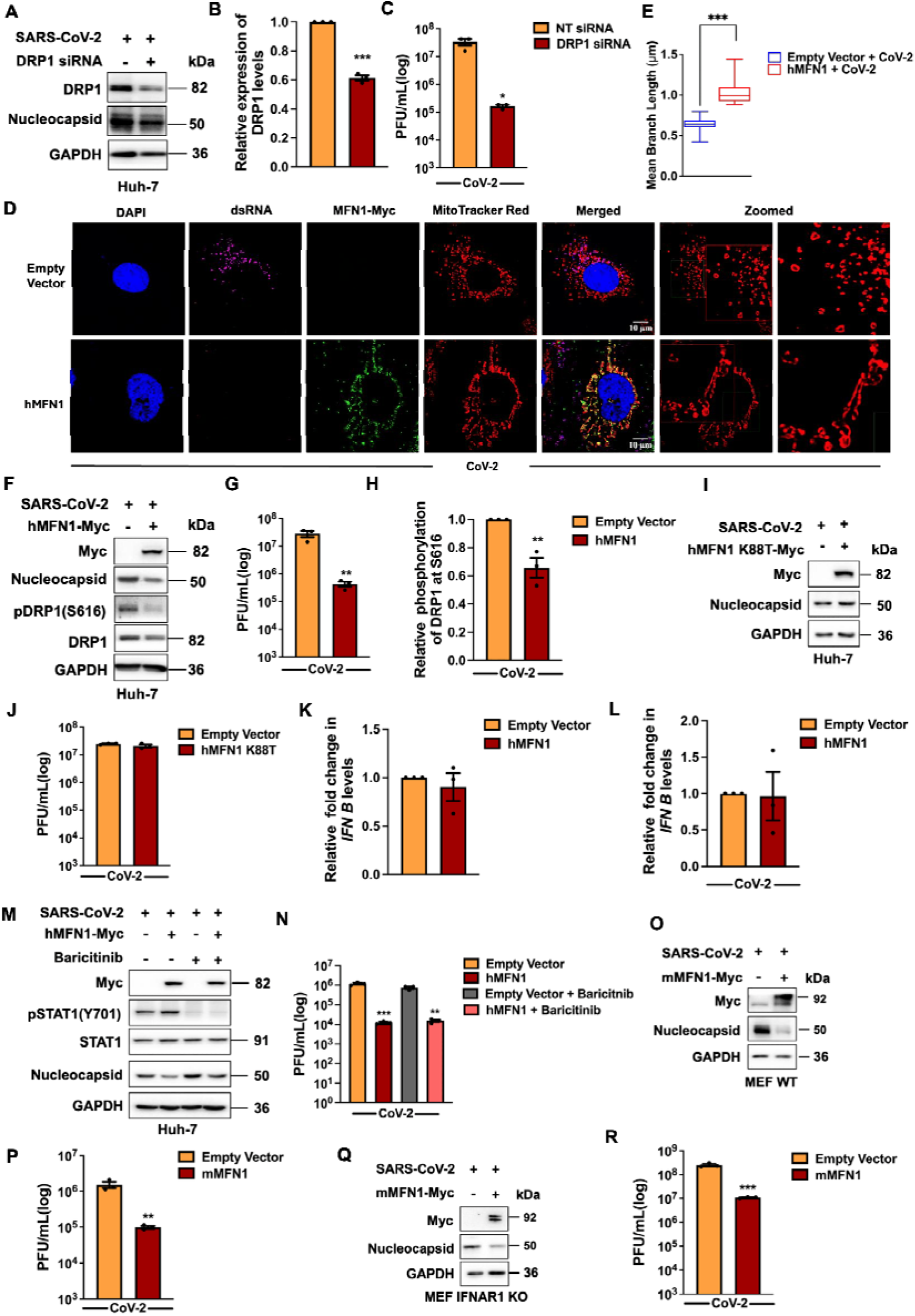
Mitochondrial fusion establishes an antiviral state in host cells independently of type I interferon signaling. (A) Immunoblots showing DRP1 and SARS-CoV-2 Nucleocapsid protein levels in Huh-7 cells following DRP1-specific siRNA transfection and SARS-CoV-2 infection. (B) Densitometric quantification of DRP1 protein levels shown in (A). (C) Infectious viral titers measured in culture supernatants from (A). (D) Confocal microscopy images showing mitochondrial elongation in SARS-CoV-2-infected Huh-7 cells overexpressing Myc-tagged human MFN1 (hMFN1-Myc), compared with empty vector controls. Mitochondria were labeled with MitoTracker Red (red). dsRNA stained (purple) and Myc-tagged hMFN1 in (green) (E) Quantification of mean mitochondrial branch length in cells shown in (D). (F) Immunoblots showing Myc-tagged human MFN1(hMFN1-Myc) expression, SARS-CoV-2 Nucleocapsid protein, pDRP1-S616 and total DRP1 levels in human MFN1-overexpressing Huh-7 cells infected with SARS-CoV-2. (G) Infectious viral titers measured in corresponding culture supernatants from (F). (H) Densitometric quantification of relative DRP1 phosphorylation at S616 shown in (F). (I) Immunoblots showing Myc-tagged GTPase-deficient human MFN1 (hMFN1-K88T-Myc) expression and SARS-CoV-2 Nucleocapsid protein levels in SARS-CoV-2 infected Huh-7 cells overexpressing the hMFN1-K88T mutant. (J) Infectious viral titers measured in culture supernatants from (I). (K) RT-qPCR analys ls. (L) RT-qPCR analysis of IFN B mRNA levels in SARS-CoV-2-infected Huh-7 cells overexpressing hMFN1- Myc compared with empty vector controls. (M) Immunoblots showing Myc-tagged human MFN1 (hMFN1-Myc) expression, phosphorylated STAT1 (pSTAT1), total STAT1, and SARS-CoV-2 Nucleocapsid protein levels in Huh-7 cells transfected with hMFN1-Myc, infected with SARS-CoV-2 and treated with baricitinib (10 µM). (N) Infectious viral titers measured in culture supernatants from (M). (O) Immunoblots showing Myc-tagged mouse MFN1 (mMFN1-Myc) expression and SARS-CoV-2 Nucleocapsid protein levels in wild-type (WT) primary MEF cells overexpressing mMFN1-Myc followed by SARS-CoV-2 infection. (P) Infectious viral titers measured in culture supernatants from (O). (Q) Immunoblots showing Myc-tagged mouse MFN1 (mMFN1-Myc) expression and SARS-CoV-2 Nucleocapsid protein levels in IFNAR1 KO primary MEF cells overexpressing mMFN1-myc followed by SARS-CoV-2 infection (R) Infectious viral titers from culture supernatants from (Q) Scale bars, 10 µm and n = 50 cells unless otherwise indicated.

Interestingly, MFN1 overexpression was also accompanied by reduced DRP1 S616 phosphorylation, suggesting attenuation of DRP1-associated fission signaling (Figure 3 F & H). Similar antiviral effects were observed following overexpression of mouse MFN1 in mouse embryonic fibroblasts (MEFs) (Figures S3 C & D). Furthermore, re-expression of MFN1 in MFN1 KO cells restored the antiviral state, strengthening the specificity of the MFN1-associated phenotype (Figures S3 E-H). We next investigated whether the antiviral activity of MFN1 extended to other RNA viruses. Interestingly, MFN1 overexpression efficiently suppressed the replication of JEV, HCV, and VSV (Figures S3 I-N), suggesting that MFN1-mediated antiviral protection is not restricted to SARS-CoV-2.

To determine whether the antiviral activity of MFN1 required its mitochondrial fusion-associated function, we examined the effects of a GTPase-deficient MFN1 mutant. Since MFN1-mediated mitochondrial fusion depends on its GTPase activity, we investigated whether the K88T MFN1 mutant retained antiviral potency. Interestingly, unlike wild-type MFN1, overexpression of the K88T mutant failed to suppress SARS-CoV-2 replication (Figures 3 I & J; Figures S3 O & P), suggesting that the antiviral activity of MFN1 is closely associated with its GTPase-dependent function. In agreement with our earlier depletion experiments, MFN2 overexpression resulted in only partial suppression of SARS-CoV-2 replication, further highlighting the comparatively stronger antiviral activity of MFN1 (Figures S3 Q & R). Collectively, these findings demonstrate that MFN1 overexpression establishes potent antiviral protection against multiple RNA viruses, with its antiviral activity closely associated with its GTPase- dependent mitochondrial remodeling function.

Previous studies have implicated MFN1-mediated redistribution of MAVS on mitochondria in promoting antiviral interferon responses^36^. However, our recent findings demonstrated that MAVS also coordinates interferon-independent intrinsic antiviral defense by preserving mitochondrial integrity in virus-infected cells^6^. We therefore investigated whether the antiviral activity of MFN1 was dependent on canonical interferon signaling. Interestingly, MFN1 overexpression did not increase basal IFNB expression in Huh-7 cells (Figure 3K). We have previously demonstrated that Huh-7 cells are competent to induce IFNB1 following poly(I:C) stimulation^6^, indicating that the absence of IFNB induction was not attributable to an intrinsic inability of these cells to mount an interferon response. We next examined IFNB expression in MFN1-overexpressing cells following SARS-CoV-2 infection. Despite potently suppressing viral replication, MFN1 overexpression did not enhance IFNB expression (Figure 3L), suggesting that its antiviral activity does not require increased type I IFN induction. Since SARS-CoV-2 is known to delay interferon responses^25,26^, we extended these observations to JEV infection, where interferon induction occurs relatively early^37^. Consistent with the SARS-CoV-2 findings, MFN1 overexpression strongly restricted JEV replication without further elevating IFNB expression compared with vector-control cells ( Figure S3S). To further investigate the involvement of interferon signaling, we treated MFN1- overexpressing, SARS-CoV-2-infected cells with baricitinib, a pharmacological inhibitor of JAK signaling, and assessed whether inhibition of this pathway reversed MFN1-mediated viral restriction. Interestingly, baricitinib treatment failed to reverse the antiviral effects of MFN1 (Figures 3 M & N), further suggesting that MFN1-associated antiviral protection does not depend on canonical IFN signaling. Finally, we examined whether MFN1 overexpression retained its antiviral activity in IFNAR1 KO MEFs that fail to induce functional type I IFN signaling. Consistent with our earlier observations, MFN1 overexpression suppressed SARS-CoV-2 replication with comparable efficiency in wild-type and IFNAR1 KO MEFs (Figures 3 O-R). Together, these findings establish that MFN1-mediated antiviral restriction operates independently of type I IFN signaling, revealing an additional contribution of MFN1-associated mitochondrial regulation to intrinsic antiviral defense.

### Inhibition of DRP1 function establishes a potent antiviral state

Following the antiviral effects associated with mitochondrial elongation in our genetic models, we next tested if inhibiting DRP1 is a potential antiviral strategy. To test this, we first treated SARS-CoV-2- infected Huh-7 cells with Mdivi-1, a widely used pharmacological inhibitor of mitochondrial fission at a concentration previously employed in Huh-7 cells^38^. Mdivi-1 treatment markedly reduced DRP1 abundance and promoted extensive mitochondrial elongation in both uninfected and SARS-CoV-2- infected Huh-7 cells compared with their untreated counterparts (Figures 4 A-D). Notably, Mdivi-1 treatment restored the mitochondrial network in infected cells to a morphology comparable to that observed in uninfected controls. Consistent with our hypothesis, Mdivi-1 treatment profoundly suppressed SARS-CoV-2 replication, as evidenced by significantly reduced viral titers (Figures 4 A & E). Mdivi-1 displayed potent antiviral activity against JEV (Figure S4A) and against SARS-CoV-2 in MEF (Figures S4 B & C), demonstrating that DRP1 inhibition promotes broad antiviral defense. Importantly, this restriction was not accompanied by increased IFNB expression (Figure S4D). In support of this, Mdivi-1 suppressed SARS-CoV-2 replication with comparable efficiency in IFNAR1 KO MEFs and their wild type counterparts (Figures S4 E & F), supporting an antiviral mechanism independent of type I IFN signaling.

**Figure 4.**
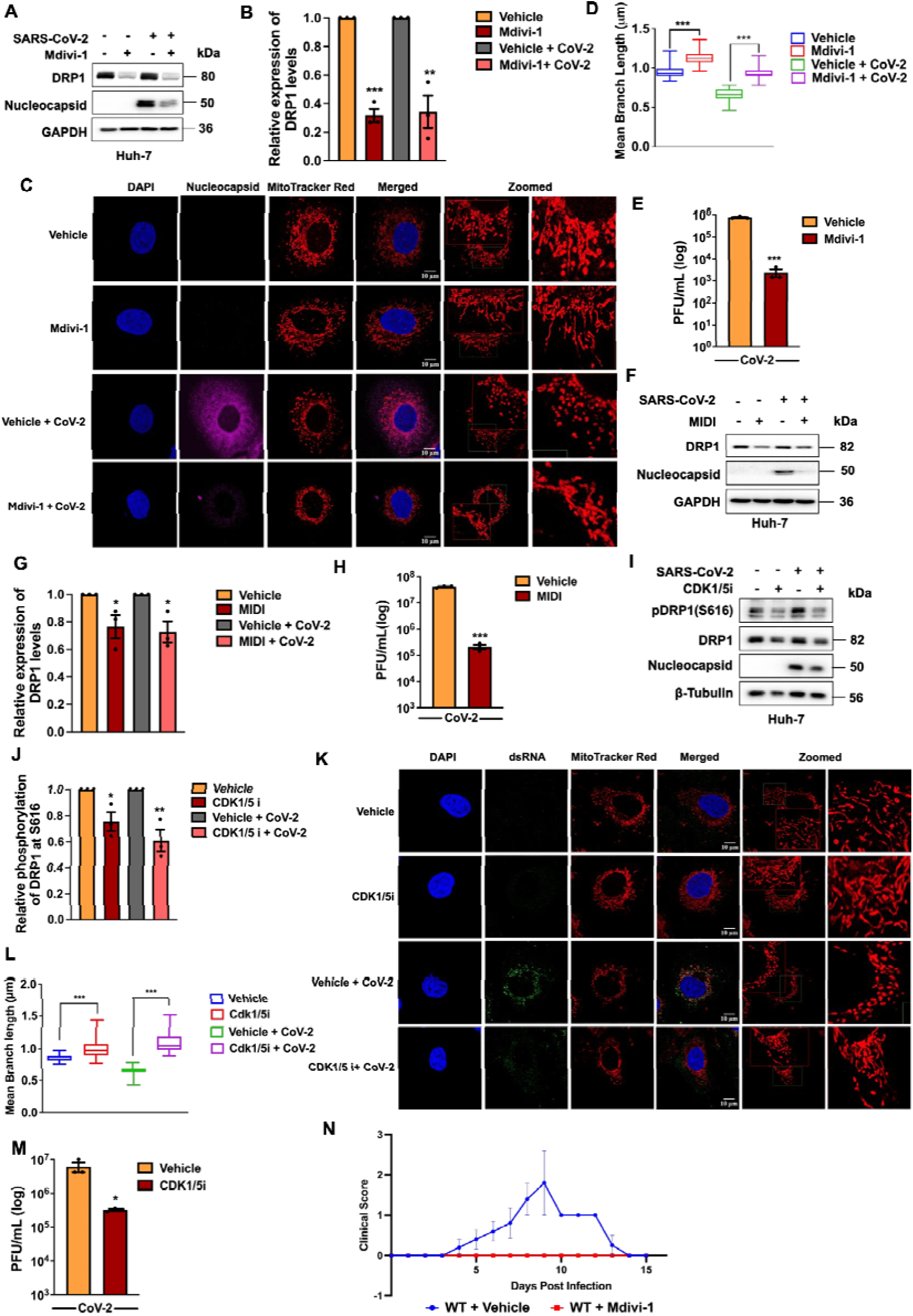
Inhibition of DRP1 function establishes a potent antiviral state. (A) Immunoblots showing DRP1 and SARS-CoV-2 Nucleocapsid protein levels in Huh-7 cells following SARS-CoV-2 infection and treatment with Mdivi-1 (50 µM). (B) Densitometric quantification of DRP1 protein levels shown in (A). (C) Confocal microscopy images showing mitochondrial morphology in uninfected and SARS-CoV-2- infected Huh-7 cells treated with Mdivi-1, compared with their respective vehicle controls. Mitochondria labelled with MitoTracker Red (red) and SARS-CoV-2 Nucleocapsid stained in (purple). (D) Quantification of mean mitochondrial branch length in cells shown in (C). (E) Infectious viral titers measured in culture supernatants from (A). (F) Immunoblots showing DRP1 and SARS-CoV-2 Nucleocapsid protein levels in SARS-CoV-2-infected Huh-7 cells treated with MIDI (5 µM). (G) Densitometric quantification of DRP1 protein levels shown in (F). (H) Infectious viral titers measured in culture supernatants from (F). (I) Immunoblots showing pDRP1-S616, total DRP1, and SARS-CoV-2 Nucleocapsid protein levels in SARS- CoV-2-infected Huh-7 cells treated with CDK1/5 inhibitor (CDK1/5i; 50 µM). (J) Densitometric quantification of relative DRP1 phosphorylation at S616 shown in (I). (K) Confocal microscopy images showing mitochondrial morphology in uninfected and SARS-CoV-2- infected Huh-7 cells treated with CDK1/5i, compared with their respective vehicle controls. (L) Quantification of mean mitochondrial branch length in cells shown in (K). (M) Infectious viral titers measured in culture supernatants from (I). (N) Clinical scores of JEV-infected mice following Mdivi-1 treatment compared with vehicle treated controls. Clinical symptoms were monitored daily from 0 to 15 days post-infection (dpi). Scale bars, 10 µm. n = 50 cells unless otherwise indicated.

Since Mdivi-1 has been reported to exert DRP1-independent effects^39^, we sought to substantiate the involvement of DRP1-associated mitochondrial fission using an alternative pharmacological approach. To this end, we employed MIDI, an inhibitor targeting the mitochondrial recruitment of DRP1 and assessed cell viability using an MTT assay which revealed 84% residual activity at 5 µM relative to controls suggesting minimal treatment associated cytotoxicity (Figure S4G). Reassuringly, MIDI treatment also markedly suppressed SARS-CoV-2 replication, with antiviral activity comparable to that observed following Mdivi-1 treatment (Figures 4 F-H). These observations further support the involvement of DRP1-associated functions in regulating viral permissiveness. We next investigated whether targeting DRP1 activation could similarly restrict viral replication. Since CDK1-mediated phosphorylation of DRP1 at S616 has been implicated in promoting its fission-associated activity^9^, we inhibited CDK1-associated signaling using CDK1/5i. Cell viability, tested using an MTT assay, revealed approximately 92% residual activity at 50μM indicating minimal treatment associated cytotoxicity (Figure S4H). Treatment with CDK1/5i efficiently reduced DRP1 S616 phosphorylation in both uninfected and SARS-CoV-2-infected cells (Figures 4 I & J). This reduction was accompanied by a marked rescue of SARS-CoV-2-induced mitochondrial fragmentation (Figures 4 K & L) and significant suppression of viral replication (Figure 4M). To further validate these findings and distinguish CDK1-associated effects from the broader activity of CDK1/5i, we performed RNAi-mediated CDK1 knockdown. Consistent with the pharmacological results, CDK1 depletion substantially reduced DRP1 S616 phosphorylation and suppressed viral replication comparable to that observed following CDK1/5i treatment (Figures 4S I-L). Together, these findings demonstrate that interventions targeting DRP1-associated mitochondrial recruitment and activation are accompanied by potent restriction of viral replication, further implicating DRP1-associated functions in the establishment of a virus-permissive cellular state.

To explore the potential relevance of these findings in vivo, we performed preliminary studies using a mouse model of JEV infection. B6129SF2/J mice were infected with JEV and subsequently treated with Mdivi-1. Interestingly, Mdivi-1 treated mice did not exhibit detectable clinical signs of infection during the 15-day observation period, whereas vehicle treated mice developed symptoms as early as 3 days post-infection (dpi), which persisted until 13 dpi (Figure 4N). Consistent with the improved in vivo outcomes, Mdivi-1-treated mice continued to gain weight throughout the observation period, whereas weight gain was markedly impaired in vehicle treated mice (Figure S4M). These observations suggest that Mdivi-1 treatment may confer protection against JEV-associated disease in vivo, extending the potential relevance of mitochondrial fission-targeted interventions beyond cultured cells.

### A major metabolic reprogramming driven by DRP1 inhibition and mitochondrial elongation supports the antiviral state

Given that mitochondria and metabolism are closely linked, we wondered whether the antiviral state associated with mitochondrial fusion involves reprogramming host-cell metabolism towards a state less permissive for viral replication. Previous studies have reported enhanced glycolytic metabolism in MFN1-deficient cells^40^. To investigate this association, we first examined the metabolic status of MFN1 KO cells. As expected, MFN1-deficient cells exhibited increased glycolytic activity, as evidenced by enhanced glycolytic flux in Seahorse glycolysis stress assays and elevated lactate levels (Figures 5 A-C). MFN1 deficiency was also accompanied by reduced succinate levels, suggesting alterations in mitochondrial oxidative metabolism (Figure 5D). SARS-CoV-2 and many RNA viruses promote glycolysis as a means to augment their replication, and inhibiting glycolysis is considered an effective strategy to restrict viral replication^16,17,41^. Therefore, we sought to establish the association between the viral restriction mediated by inhibition of glycolysis and alterations in mitochondrial dynamics. Treatment with the glycolytic inhibitors 2-deoxyglucose (2-DG) and DL-glyceraldehyde (DL-G) markedly suppressed SARS-CoV-2 replication (Figures 5 E-H; Figures S5 A & B) while simultaneously promoting extensive mitochondrial elongation (Figures 5 I & J). To determine whether mitochondrial elongation occurred independently of viral restriction, we examined mitochondrial morphology in uninfected cells following glycolytic inhibition. Notably, both inhibitors induced pronounced mitochondrial elongation even in the absence of infection (Figures S5 C & D), indicating that the morphological changes were induced by glycolytic inhibition rather than arising solely as a consequence of reduced viral replication.

**Figure 5.**
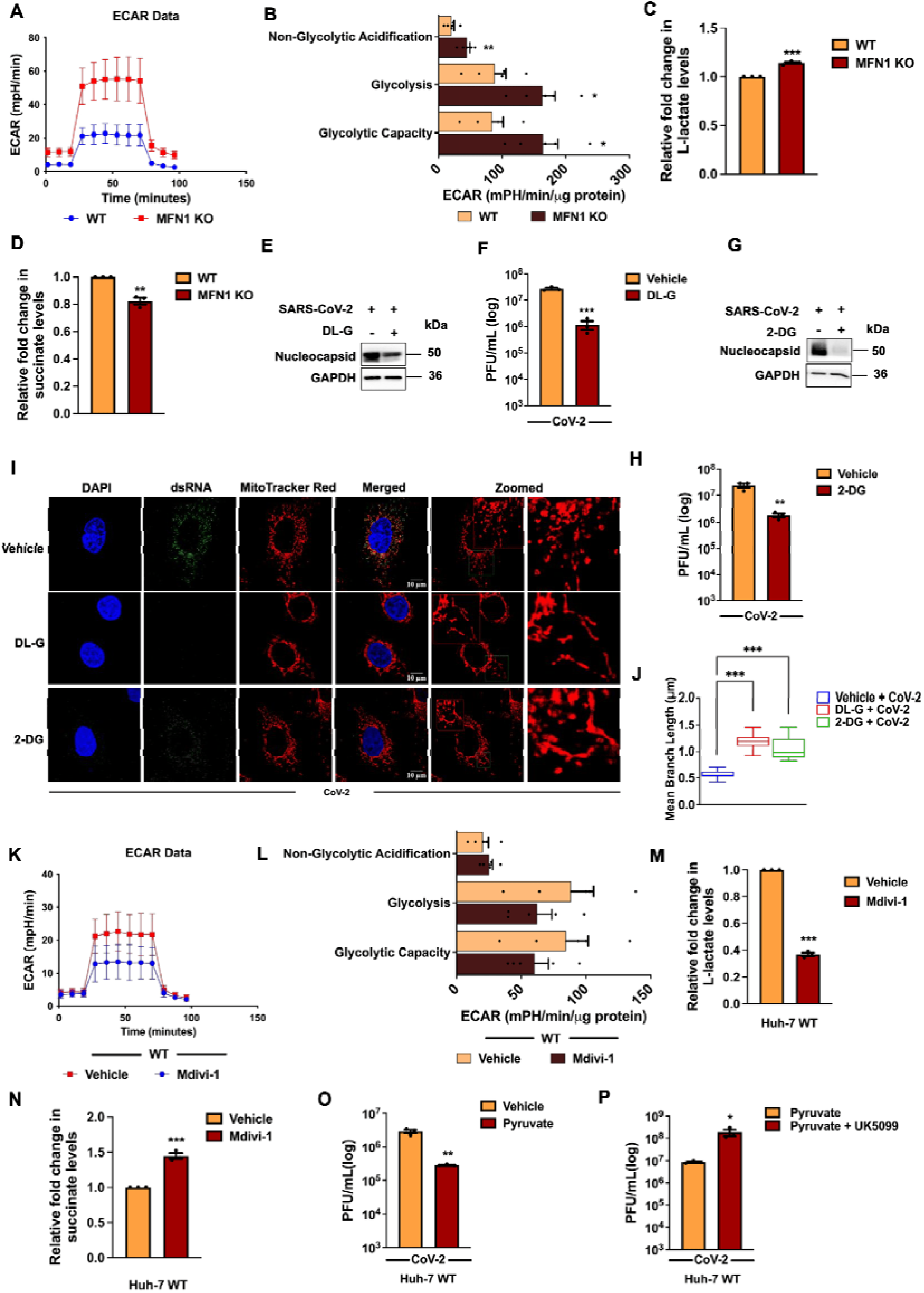
A major metabolic reprogramming driven by DRP1 inhibition and mitochondrial elongation supports the antiviral state. (A) Extracellular acidification rate (ECAR) profiles and (B) glycolytic parameters in wild-type (WT) and MFN1 KO Huh-7 cells measured using the Seahorse XFe24 Glycolysis Stress Test (n = 5). (C) L-lactate levels measured in culture supernatants from WT and MFN1 KO Huh-7 cells. (D) Intracellular succinate levels measured in WT and MFN1 KO Huh-7 cells. (E) Immunoblots showing SARS-CoV-2 Nucleocapsid protein levels in Huh-7 cells following SARS-CoV-2 infection and treatment with DL-G. (F) Infectious viral titers measured in culture supernatants from (E). (G) Immunoblots showing SARS-CoV-2 Nucleocapsid protein levels in SARS-CoV-2-infected Huh-7 cells treated with 2-DG. (H) Infectious viral titers measured in culture supernatants from (G). (I) Confocal microscopy images showing mitochondrial morphology in SARS-CoV-2-infected Huh-7 cells treated with DL-G or 2-DG, compared with vehicle control. (J) Quantification of mean mitochondrial branch length in cells shown in (I). (K) ECAR profiles and (L) glycolytic parameters in vehic le- and Mdivi-1-treated Huh-7 cells measured using the Seahorse XFe24 Glycolysis Stress Test (n = 5). (M) L-lactate levels measured in culture supernatants from vehicle- and Mdivi-1-treated Huh-7 cells. (N) Intracellular succinate levels measured in vehicle- and Mdivi-1-treated Huh-7 cells. (O) Infectious viral titers measured in culture supernatants from SARS-CoV-2-infected Huh-7 cells supplemented with sodium pyruvate (440 mg/L), compared with vehicle controls. (P) Infectious viral titers measured in culture supernatants from SARS-CoV-2-infected Huh-7 cells supplemented with sodium pyruvate (440 mg/L) and treated with UK5099 (25 µM). Scale bars, 10 µm. n = 50 cells unless otherwise indicated.

We next investigated the possibility of metabolic reprogramming behind Mdivi-1’s antiviral functions. Interestingly, Mdivi-1 markedly suppressed glycolytic metabolism in Huh-7 cells, as evidenced by reduced glycolytic flux in Seahorse glycolysis stress assays and decreased lactate levels (Figures 5 K-M). Furthermore, Mdivi-1 treatment increased succinate abundance (Figure 5N), suggesting a shift away from glycolytic metabolism accompanied by alterations in mitochondrial carbon metabolism. To further investigate the potential contribution of mitochondrial oxidative metabolism to antiviral restriction, we supplemented SARS-CoV-2-infected Huh-7 cells with sodium pyruvate and examined its effects on mitochondrial morphology and viral replication. Pyruvate treatment markedly restored mitochondrial elongation in infected cells while simultaneously suppressing SARS-CoV-2 replication (Figure 5O; Figures S5 E & F). Interestingly, inhibition of mitochondrial pyruvate import using the mitochondrial pyruvate carrier inhibitor UK5099 reversed the antiviral effect of pyruvate supplementation (Figure 5P), suggesting that mitochondrial pyruvate uptake contributes to the observed viral restriction. Cell viability assessment using an MTT assay revealed approximately 95% residual activity of UK5099 (25 µM), suggesting minimal treatment-associated cytotoxicity (Figure S5G). These results collectively indicated that the antiviral state established by robust mitochondrial fusion mediated by DRP1 inhibition is orchestrated through lowering glycolysis flux and by promoting mitochondrial oxidative metabolic state.

### MFN1 and mitochondrial fusion are dispensable for DRP1-mediated viral permissiveness

Although our findings so far established strong associations between mitochondrial fusion and potent antiviral state, and between mitochondrial fragmentation and enhanced viral permissiveness, whether mitochondrial dynamics itself regulates the viral permissiveness, or whether it is regulated independently by DRP1 and MFN1 remained unclear. We therefore investigated whether the antiviral effects of DRP1 inhibition depended on MFN1-mediated mitochondrial fusion. Surprisingly, Mdivi-1 treatment efficiently suppressed SARS-CoV-2 replication in MFN1 KO cells, with antiviral activity comparable to that observed in wild-type (WT) cells (Figures 6 A & B). Confocal imaging revealed that, unlike the pronounced mitochondrial elongation observed in WT cells following Mdivi-1 treatment (Figures 4 C & D), mitochondria remained extensively fragmented in MFN1 KO cells despite robust viral suppression (Figures 6 C & D). Similarly, MIDI treatment effectively restricted SARS-CoV-2 replication in MFN1 KO cells (Figures 6 E & F). These findings provide two important insights. First, mitochondrial elongation is not essential for the antiviral activity associated with DRP1-targeted interventions. Second, MFN1 itself is dispensable for this antiviral restriction, suggesting that DRP1-associated functions regulate viral permissiveness independently of MFN1-mediated mitochondrial fusion.

**Figure 6.**
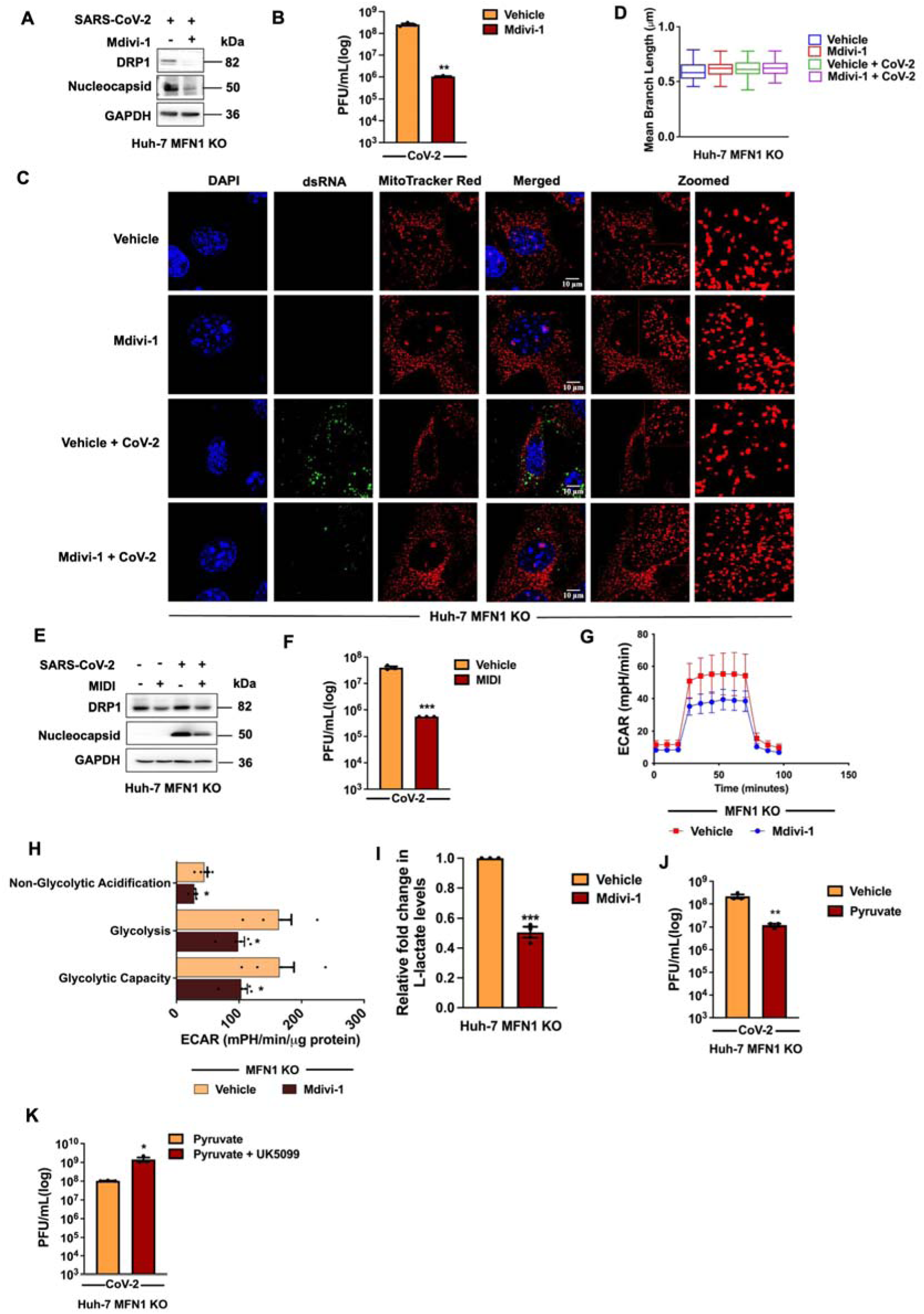
MFN1 and mitochondrial fusion are dispensable for DRP1 mediated viral permissiveness. (A) Immunoblots showing DRP1 and SARS-CoV-2 Nucleocapsid protein levels in SARS-CoV-2-infected MFN1 KO Huh-7 cells treated with Mdivi-1. (B) Infectious viral titers measured in culture supernatants from (A). (C) Confocal microscopy images showing mitochondrial morphology in uninfected and SARS-CoV-2- infected MFN1 KO Huh-7 cells treated with Mdivi-1, compared with their respective vehicle controls. (D) Quantification of mean mitochondrial branch length in cells shown in (C). (E) Immunoblots showing DRP1 and SARS-CoV-2 Nucleocapsid protein levels in SARS-CoV-2-infected MFN1 KO Huh-7 cells treated with MIDI (5 µM). (F) Infectious viral titers measured in culture supernatants from (E). (G) Extracellular acidification rate (ECAR) profiles and (H) glycolytic parameters in vehicle- and M divi-1- treated MFN1 KO Huh-7 cells measured using the Seahorse XFe24 Glycolysis Stress Test (n = 5). (I) L-lactate levels measured in culture supernatants from vehicle- and Mdivi-1-treated MFN1 KO Huh-7 cells. (J) Infectious viral titers measured in culture supernatants from SARS-CoV-2-infected MFN1 KO Huh-7 cells supplemented with sodium pyruvate (440 mg/L), compared with vehicle controls. (K) Infectious viral titers measured in culture supernatants from SARS-CoV-2-infected MFN1 KO Huh-7 cells supplemented with sodium pyruvate (440 mg/L) and treated with UK5099 (25 µM). Scale bars, 10 µm. n = 50 cells unless otherwise indicated.

Given that Mdivi-1 retained comparable antiviral activity in WT and MFN1 KO cells, we next investigated whether MFN1 is dispensable for its antiviral effects and metabolic reprogramming associated with it.

Consistent with our observations in WT cells (Figures 5 K-M), Mdivi-1 treatment markedly suppressed glycolytic metabolism in MFN1 KO cells, as evidenced by reduced glycolytic flux in Seahorse glycolysis stress assays and decreased lactate levels (Figures 6 G-I). This was accompanied by increased succinate levels (Figure S6A), suggesting alterations in mitochondrial oxidative metabolism alongside glycolytic suppression. We next verified whether pyruvate supplementation could similarly restrict viral replication in MFN1 KO cells. Interestingly, pyruvate treatment significantly suppressed SARS-CoV-2 replication (Figure 6J) without substantially altering mitochondrial morphology (Figures S6 B & C), ruling out the association between mitochondrial elongation and a robust antiviral state. Moreover, as observed in WT cells, inhibition of mitochondrial pyruvate import using UK5099 reversed the antiviral activity of exogenous pyruvate, resulting in increased viral titers compared with pyruvate treatment alone (Figure 6K). These sets of results collectively demonstrate that MFN1 and its associated mitochondrial elongation are dispensable in the regulatory oversight of glycolysis and mitochondrial metabolism underlying the antiviral state in host cells.

DRP1-mediated viral permissiveness encompasses broader mechanisms beyond glycolytic metabolism Interestingly, we observed that treatment with the glycolytic inhibitors 2-deoxyglucose (2-DG) and DL- glyceraldehyde (DL-G) resulted in a marked reduction in DRP1 abundance (Figures 7 A-D), revealing an association between glycolytic inhibition and reduced DRP1 protein levels. This observation complemented our earlier findings that interventions affecting DRP1 abundance, phosphorylation, or mitochondrial recruitment were accompanied by potent suppression of SARS-CoV-2 replication (Figures 3 A-C and 4 A-M; Figures S4 G-J). We therefore investigated whether direct depletion of DRP1 could restrict viral replication independently of MFN1. To address this, we performed shRNA-mediated DRP1 knockdown in both WT and MFN1 KO cells. Interestingly, DRP1 depletion resulted in robust and comparable suppression of SARS-CoV-2 replication in both cellular backgrounds (Figures 7 E-J), further establishing that the antiviral effects associated with loss of DRP1 do not require MFN1-dependent mitochondrial elongation. Importantly, DRP1 depletion or targeting resulted in substantially greater viral suppression than direct glycolytic inhibition, suggesting that DRP1-associated functions may extend beyond the regulation of glycolysis in determining viral permissiveness. Given this difference in antiviral activity, together with the reduction in DRP1 abundance following glycolytic inhibition, we next investigated whether simultaneous targeting of DRP1 and glycolysis could further enhance viral restriction. Interestingly, combined treatment with MIDI and DL-G resulted in only a modest additional reduction in SARS-CoV-2 titers compared with MIDI alone (Figure 7K), without the pronounced enhancement in viral restriction anticipated from dual inhibition, suggesting that glycolytic regulation may represent one component of the broader cellular processes associated with DRP1-dependent viral permissiveness.

**Figure 7.**
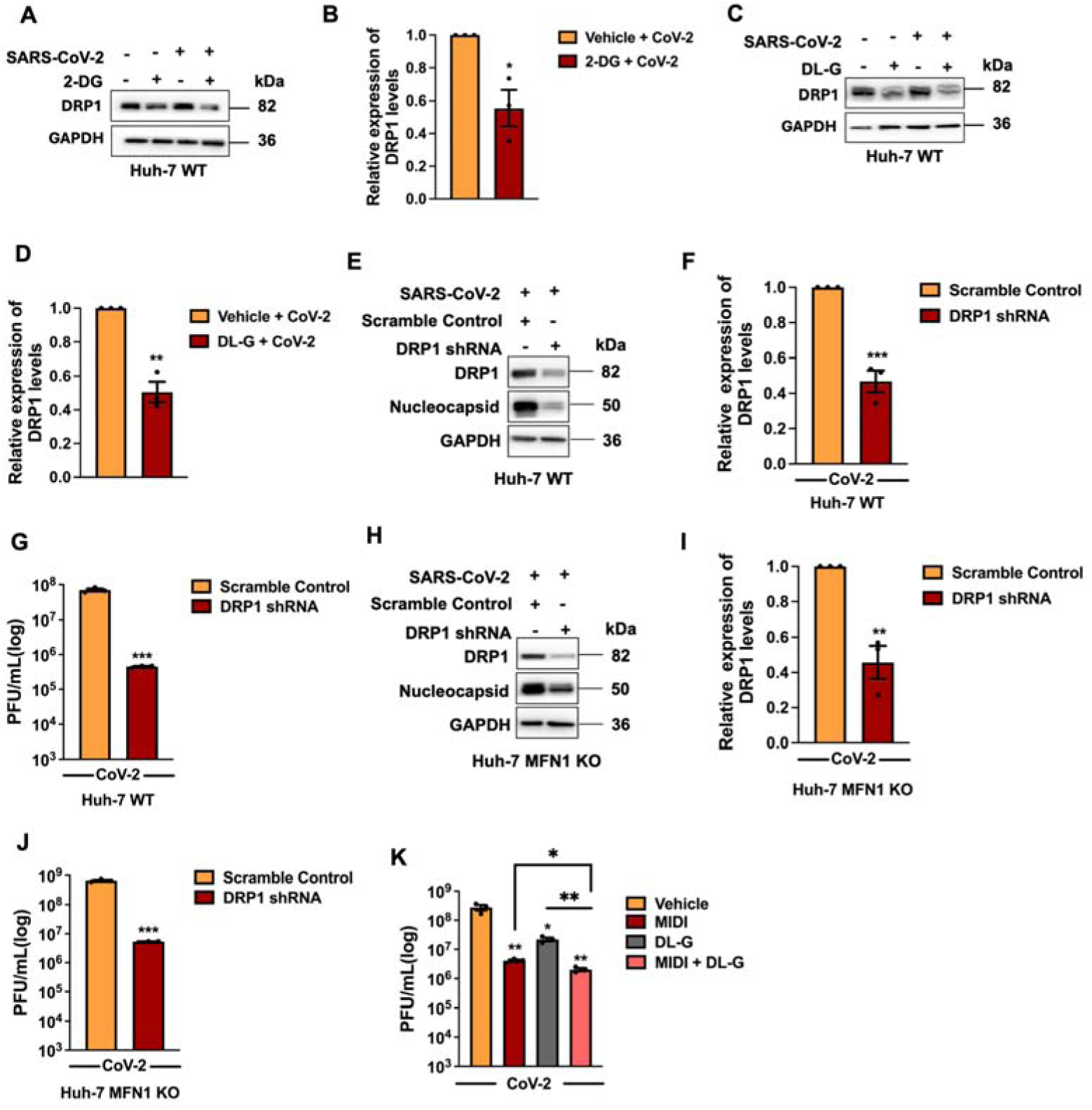
DRP1-mediated viral permissiveness encompasses broader mechanisms beyond glycolytic metabolism (A) Immunoblots showing DRP1 protein levels in uninfected and SARS-CoV-2-infected Huh-7 cells treated with 2-deoxyglucose (2-DG). (B) Densitometric quantification of DRP1 protein levels shown in (A). (C) Immunoblots showing DRP1 protein levels in uninfected and SARS-CoV-2-infected Huh-7 cells treated with DL-glyceraldehyde (DL-G). (D) Densitometric quantification of DRP1 protein levels shown in (C). (E) Immunoblots showing DRP1 and SARS-CoV-2 Nucleocapsid protein levels in Huh-7 cells co- transfected with two DRP1-targeting shRNAs (shRNA1 and shRNA2), compared with scrambled shRNA controls, following SARS-CoV-2 infection. (F) Densitometric quantification of DRP1 protein levels shown in (E). (G) Infectious viral titers measured in culture supernatants from (E) . (H) Immunoblots showing DRP1 and SARS-CoV-2 Nucleocapsid protein levels in MFN1-KO Huh-7 cells co- transfected with two DRP1-targeting shRNAs (shRNA1 and shRNA2), compared with scrambled shRNA controls, following SARS-CoV-2 infection. (I) Densitometric quantification of DRP1 protein levels shown in (H). (J) Infectious viral titers measured in culture supernatants from (H). (K) Infectious viral titers measured in culture supernatants from SARS-CoV-2-infected Huh-7 cells treated with MIDI, DL-G, or their combination (MIDI + DL-G), compared with vehicle controls.

Collectively, these findings reveal an important distinction between mitochondrial morphology and the cellular functions associated with DRP1 in regulating viral permissiveness. Although MFN1 overexpression establishes a potent antiviral state, neither MFN1 nor the mitochondrial elongation it promotes is required for antiviral restriction. Instead, inhibition of DRP1 establishes a potent antiviral state irrespective of MFN1, accompanied by suppression of glycolytic metabolism. Furthermore, the greater antiviral effect of DRP1 targeting compared with direct glycolytic inhibition suggests that DRP1-associated functions extend beyond glycolytic regulation. Together, these results identify DRP1 as the central determinant of the virus-permissive cellular state.

## DISCUSSION

Viruses exploit the metabolic plasticity of host cells to create an intracellular environment favorable for their replication. Mitochondria occupy a central position in this process, coordinating energy production, metabolic adaptation, organelle homeostasis, and antiviral immune signaling^42,43^. Their dynamic organization is therefore increasingly recognized as an important component of virus–host interactions. However, whether mitochondrial remodeling directly governs viral replication or reflects broader changes in cellular metabolic activity remains poorly understood. Two important insights emerge from our study. First, mitochondrial dynamics associated proteins influence the metabolic state determining viral permissiveness, with MFN1 deficiency favoring a glycolytic, virus-permissive environment. Second, and more importantly, mitochondrial elongation itself is not essential for establishing an antiviral state. Instead, DRP1 emerges as a central determinant of viral permissiveness, whose influence on viral replication extends beyond its conventional role in mitochondrial fission and its association with glycolytic metabolism. Our findings therefore distinguish the regulatory functions of mitochondrial dynamics-associated proteins from the morphological changes they coordinate, highlighting their broader contributions to the cellular environment supporting viral replication.

The ability of viruses to manipulate mitochondrial dynamics is increasingly recognized as an important aspect of their interaction with host cells. Mitochondrial fragmentation has frequently been associated with viral infection, yet its functional significance in determining viral permissiveness remains poorly understood. In our study, SARS-CoV-2 promoted early activation of DRP1-associated mitochondrial fission, followed by mitochondrial fragmentation, increased mitophagy, and eventual loss of mitochondrial mass. Similar fragmentation during JEV, HCV, and VSV infections suggests that mitochondrial remodeling is a recurring feature of RNA-virus infection. Interestingly, contrasting observations have also been reported, including SARS-CoV-2-induced mitochondrial elongation and enhanced oxidative metabolism supporting viral propagation^44^. These differences may arise from variations in cellular backgrounds, infection stages, and experimental conditions, highlighting the complexity of the relationship between mitochondrial morphology and viral replication. In our experimental models, MFN1 deficiency markedly enhanced viral replication, whereas MFN1 overexpression established potent antiviral protection against multiple RNA viruses. Previous studies have implicated MFN1 mediated mitochondrial fusion in promoting MAVS-dependent interferon signaling^36,45^. However, its antiviral activity in our study persisted independently of type I interferon responses. The delayed interferon induction associated with SARS-CoV-2 infection provided an opportunity to investigate this intrinsic antiviral activity, which was further supported using JEV infection and IFNAR1-deficient cells. These observations complement our recent findings demonstrating that MAVS-mediated preservation of mitochondrial integrity contributes to interferon-independent antiviral defense^6^. An important question arising from these observations is how mitochondrial dynamics- associated proteins influence the cellular metabolic environment supporting viral replication. Enhanced glycolysis has been widely implicated in promoting RNA-virus replication, including HIF-1α-associated glycolytic reprogramming during SARS-CoV-2 infection^41^. Our findings extend these observations by demonstrating that MFN1 deficiency favors a glycolytic, virus-permissive environment, whereas interventions targeting DRP1-associated functions suppress glycolytic metabolism and viral replication. Interestingly, glycolytic inhibition also promoted mitochondrial elongation, further supporting an association between mitochondrial remodeling and metabolic regulation. Together, these observations raise a key question as to whether the accompanying morphological changes are essential for establishing the antiviral state.

A particularly important implication of our findings is that mitochondrial elongation, despite its strong association with antiviral protection, is not necessarily the underlying requirement for viral restriction. This became evident when inhibition of DRP1-associated functions retained potent antiviral activity in MFN1-deficient cells despite persistent mitochondrial fragmentation. More importantly, glycolytic suppression and pyruvate-mediated antiviral restriction remained effective under these conditions, demonstrating that the metabolic processes associated with viral restriction can operate independently of MFN1-mediated mitochondrial elongation. These findings challenge the conventional interpretation of mitochondrial morphology as a direct indicator of cellular metabolic competence. Although mitochondrial fragmentation is frequently associated with metabolic dysfunction and mitochondrial elongation with improved organelle function, the relationship between structure and function is evidently more complex^15,46,47^. Our observations suggest that mitochondrial remodeling may accompany the establishment of particular metabolic states without necessarily being their determining factor. In this context, the antiviral benefit associated with mitochondrial fusion may arise from the metabolic and cellular functions it supports rather than from mitochondrial elongation itself. This distinction is important for understanding virus-induced mitochondrial remodeling, as it suggests that the functional consequences of altered mitochondrial dynamics cannot be inferred solely from changes in mitochondrial architecture.

The separation between mitochondrial morphology and antiviral activity identifies DRP1 as an important regulatory component of the virus-permissive cellular state. DRP1 is traditionally recognized as a central regulator of mitochondrial fission, yet emerging evidence indicates that its cellular functions are closely associated with metabolic adaptation. Nagdas and colleagues demonstrated that DRP1 supports oncogenic KRAS-driven glycolysis while also maintaining mitochondrial metabolic functions through partially independent mechanisms^48^. Our findings extend this relationship to viral infection, where interventions affecting DRP1 abundance, activation, or mitochondrial recruitment were associated with potent antiviral restriction. Surprisingly, glycolytic inhibition markedly reduced DRP1 abundance, while Mdivi-1 treatment suppressed glycolytic activity, suggesting functional interplay between the mitochondrial fission machinery and cellular metabolism. The importance of DRP1 was further supported by genetic depletion, which strongly restricted viral replication even in the absence of MFN1. Consistently, a strong DRP1 dephosphorylation was observed during MFN1 overexpression, implicating the attenuation of DRP1-associated activity behind the potent antiviral protection conferred by MFN1. This observation provides a possible functional connection between MFN1-mediated antiviral protection and DRP1 targeting, although the extent to which reduced DRP1 activation accounts for the antiviral effects of MFN1 remains to be determined. Notably, the magnitude of viral suppression following DRP1 targeting substantially exceeded that achieved by glycolytic inhibition alone. This raises the possibility that DRP1 contributes to viral permissiveness through additional functions beyond glycolytic regulation. Given the involvement of mitochondrial dynamics in organelle quality control, metabolic adaptation, and intraorganellar communication, DRP1-associated cellular processes may collectively influence the availability of metabolic resources and the maintenance of an intracellular environment favorable for viral replication.

The possibility of broader DRP1-associated metabolic functions is further supported by its extensive protein interaction network. Our analysis of BioGRID (Supplementary Figure 7) revealed associations between DRP1 and proteins involved in several metabolic pathways, including glycolysis and the pentose phosphate pathway (ALDOA, LDHA, TKT), nucleotide biosynthesis (RRM1, RRM2, IMPDH2), mitochondrial carbon metabolism (PDHB, MDH2, SUCLG2), lipid metabolism (ACLY, FASN), and redox homeostasis (PRDX2, SOD1, TXN)^49^. Interestingly, these metabolic processes collectively represent major cellular requirements for viral replication, providing the energy, nucleotides, membrane components, and redox balance necessary for efficient viral propagation. The distribution of DRP1-associated proteins across these pathways raises the possibility that DRP1 participates in a broader cellular network capable of influencing metabolic permissiveness beyond glycolysis alone. Although protein associations do not necessarily indicate direct physical interactions or regulatory control, this network offers a framework for understanding how perturbation of DRP1-associated functions could influence multiple cellular processes simultaneously. These associations may help explain why the antiviral effects of DRP1 targeting are more pronounced than those achieved through glycolytic inhibition alone, although contributions from nonmetabolic processes cannot be excluded. Thus, rather than viewing DRP1 solely as an executor of mitochondrial division, our findings encourage consideration of its broader association with cellular metabolic homeostasis and the metabolic requirements of viral replication.

These findings also raise important questions regarding the therapeutic potential of targeting DRP1- associated cellular functions. Host-directed antiviral approaches offer an opportunity to disrupt cellular processes required for viral replication, potentially providing protection against viruses that share common metabolic dependencies. The broad antiviral effects observed following MFN1 overexpression and interventions targeting DRP1-associated regulation suggest that mitochondrial metabolic pathways may represent useful points of therapeutic intervention. Preliminary observations from our JEV mouse model further support the potential relevance of these findings beyond cultured cells. However, DRP1 performs essential physiological functions, and broadly disrupting mitochondrial fission may have consequences for mitochondrial quality control and cellular homeostasis. Moreover, Mdivi-1 has been reported to inhibit mitochondrial complex I independently of DRP1^39^, indicating that its antiviral and metabolic effects cannot be attributed exclusively to DRP1 inhibition. Although our genetic depletion experiments independently establish a contribution of DRP1 to viral permissiveness, the extent to which its antiviral effects are mediated by altered glycolysis, mitochondrial metabolism, or additional cellular processes remains unresolved. Further studies combining genetic DRP1 perturbation with metabolic flux analysis and targeted interrogation of its associated metabolic pathways will therefore be important for identifying the functions responsible for viral restriction. Such approaches may reveal opportunities to disrupt virus-supportive metabolic pathways without compromising the essential physiological functions of mitochondrial fission.

Together, our findings suggest that the relationship between mitochondrial dynamics and viral replication extends beyond the conventional opposition between mitochondrial fusion and fission. While MFN1-associated mitochondrial elongation accompanies potent antiviral activity, the persistence of viral restriction in cells with fragmented mitochondria demonstrates that mitochondrial architecture alone does not determine the antiviral state. Instead, our study places DRP1 at the interface of mitochondrial dynamics, metabolic regulation, and viral permissiveness, with its contribution to viral replication potentially extending across multiple cellular processes. This distinction broadens our understanding of how mitochondrial-associated functions influence host susceptibility to infection and suggests that disruption of the metabolic environment supporting viral replication may be more consequential than restoration of mitochondrial morphology itself. By identifying DRP1 as a central determinant of viral permissiveness, our findings provide a framework for investigating how mitochondrial dynamics-associated metabolic networks contribute to intrinsic antiviral defense and offer potential avenues for host-directed antiviral intervention.

## LIMITATIONS OF THE STUDY

Our findings identify DRP1 as an important regulator of RNA virus permissiveness, yet several questions remain unanswered. The molecular mechanisms linking DRP1 to metabolic reprogramming independently of mitochondrial morphology remain to be elucidated. While pharmacological inhibition and genetic depletion consistently restrict viral replication, the relative contributions of glycolytic suppression and other potential DRP1-associated cellular functions require further investigation.

Additionally, changes in glycolytic activity and metabolite abundance suggest metabolic reprogramming, but direct measurements of mitochondrial oxidative metabolism are necessary to establish its functional contribution. Finally, complementary pharmacological and genetic approaches strengthen our conclusions, but further mechanistic investigations and validation in additional physiological and in vivo models will be essential to determine the broader relevance and therapeutic potential of targeting DRP1-associated pathways.

## MATERIALS AND METHODS

### Antibodies

Phospho DRP1 S616 (Cat #3455), DRP1 (Cat #8570), Mitofusin 1 (MFN1) reactive to human (Cat #14739), Mitofusin 2 (MFN2) (Cat #9482), MFF (Cat #84580), CDC2 (CDK1) (Cat #77055), Myc tag (Cat #2278), GFP tag (Cat #2956), phospho-STAT1 (Cat #14994) and STAT1 (Cat #9167), were bought from Cell Signaling Technology (CST). JEV NS1 (Cat #GT1410) antibody was from Genetex. dsRNA J2 antibody (Cat #10010200) was procured from Scicons. Mitofusin 1(MFN1) (Cat #ab126575) reactive to mouse and HCV NS3 (Cat #ab21124) was bought from Abcam. Phospho PINK1 S228 (Cat #PA5-105356), PINK1 (Cat #PA5- 86941), CoV-2 Nucleocapsid (Cat #MA5-36272), GAPDH (Cat #MA5-15738) were procured from Invitrogen. Goat anti-rabbit HRP conjugated (Cat #115035-146), Goat anti-mouse HRP conjugated (Cat #115035-003), Alexa Fluor 488 conjugated (Cat #115-545-003), Alexa Fluor 594 conjugated (Cat #111-585-006) and Alexa Fluor 647 conjugated (Cat #115-605-003) were purchased from Jakson ImmunoResearch.

### Plasmids and cloning

Mammalian expression vector, pcDNA4/TO, was from Invitrogen. Human MFN1-Myc, MFN2 and MFF were amplified using Huh-7 cDNA and cloned into pcDNA4/TO. Human MFN1 K88T mutation was generated by overlap SDM and cloned into pcDNA4/TO. Human DRP1 pEGFPC1 plasmid was kindly provided by Dr. Michael T. Ryan ^50^. Mouse MFN1 (Plasmid #23212) and MFN1 K88T (Plasmid #26050) pcDNA3.1Myc plasmids were procured from Addgene.

### siRNA and shRNA transfections

siRNAs (AM16708) targeting human DRP1(Assay ID:146450), MFN1(Assay ID:138495), MFN2 (Assay ID:133649), and CDC2(CDK1) (Assay ID:103821), and NT siRNA are procured from Ambion (Invitrogen). 5nmoles of indicated siRNA oligos were transfected to cells using Oligofectamine 2000 according to manufacturer instructions. shRNA scramble pLKO.1 plasmid was procured from Addgene (Plasmid #162011). shRNA oligos targeting DRP1 were annealed and cloned into pLKO.1 TRC. These shRNA’s were transfected to cells using lipofectamine 3000 reagent according to manufacturer instructions.

### Real-time quantitative PCR (RT-qPCR)

Cellular RNA samples were prepared using an MN NucleoSpin RNA kit (Macherey-Nagel, Cat #740955.50). Equal quantities of RNA were reverse-transcribed using Primescript 1st-strand cDNA synthesis kit (TaKaRa, Cat #6110) following the manufacturer’s protocol. Fifty nanograms of cDNA was used for quantification using TB Green Premix (TaKaRa, Cat #RR820) on a LightCycler 480 instrument (Roche). Transcripts of the host origin were normalized against GAPDH. Relative fold changes between the experimental and control samples (2−ΔΔCT) were calculated and represented in the graphs. Table 2 details the primer sequences used for the amplification.

**TABLE 1.**
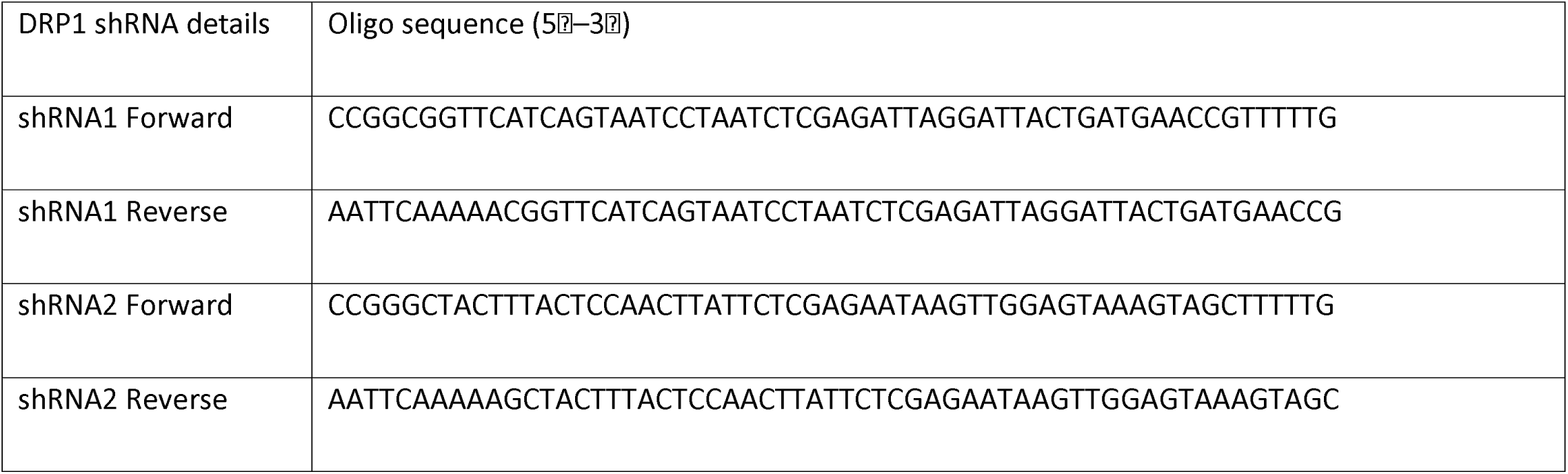
DRP1 shRNA oligos

**TABLE 2.**
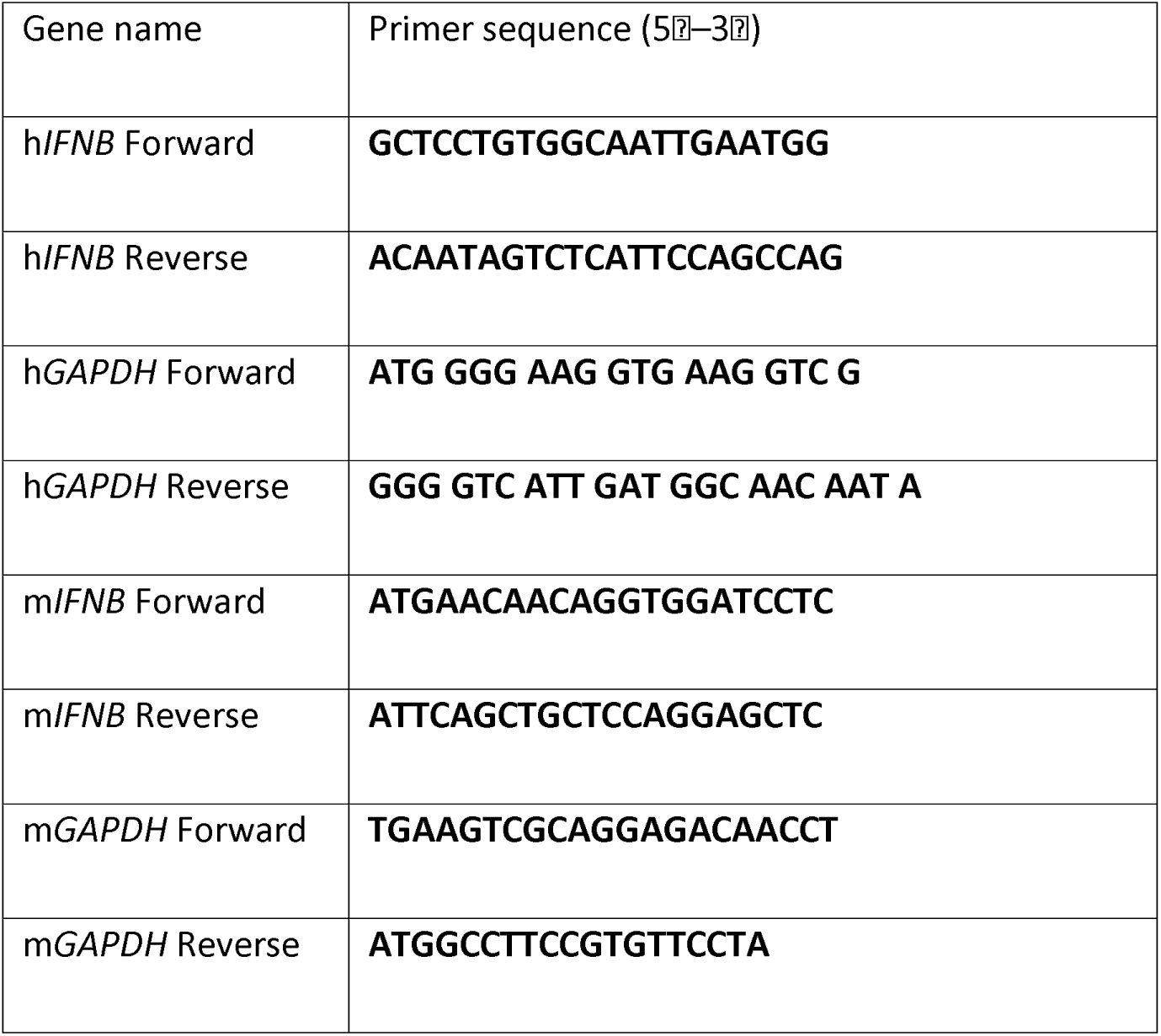
RT primers

### Cell culture

Huh-7 cells were kindly provided by Ralf Bartenschlager ^51^. Immortalized wild-type (WT) and MFN1 knockout (KO) mouse embryonic fibroblasts (MEFs) were obtained from ATCC (CRL-2991 and CRL-2992). Primary WT and IFNAR1 KO MEFs were isolated from WT and IFNAR1 KO hybrid mice generated by crossing B6129SF2/J and C57BL/6J strains, following a previously established protocol^52^. All cell lines were cultured in Dulbecco’s modified Eagle’s medium (DMEM; Gibco) supplemented with 10% fetal bovine serum (FBS; Gibco), 1× penicillin–streptomycin (Gibco), and 1× non-essential amino acids (Gibco). All cells were maintained at 37°C in a humidified incubator containing 5% CO₂. Cells were routinely passaged upon reaching approximately 70–80% confluency and were periodically screened for mycoplasma contamination.

### SARS-CoV-2, JEV, HCV and VSV isolates

The B.1.1.8 (A2a) variant (GISAID ID - EPI_ISL_458046) of SARS-CoV-2 used in this study was isolated at the Centre for Cellular and Molecular Biology in the biosafety level 3 facility^53^. The viruses were propagated in Vero (CCL-81) cells grown in serum-free media (SFM). JEV, HCV and VSV were propagated as previously described^54^.

### Virus infection, treatments, and quantification

Cells were infected with SARS-CoV-2 or JEV at a multiplicity of infection (MOI) of 1 for 3 h, VSV at an MOI of 0.5 for 1 h, or HCV at an MOI of 1 for 4 h, under serum-free conditions. The inoculum was then replaced with complete medium, and cells were incubated until the designated harvest time. At harvest, cells were trypsinized and collected separately for protein and RNA analyses. SARS-CoV-2, JEV, and VSV titers were determined by plaque assay as previously described^53,54^. HCV titers were determined by RT- qPCR as described previously^54^. For overexpression experiments, cells were seeded to achieve 50%–60% confluency and transfected with 2μg plasmid of interest using Opti-MEM and Lipofectamine 3000 according to the manufacturer’s instructions. After 6 h, the transfection mixture was replaced with complete DMEM, and cells were incubated for an additional 18 h. At 24 h post-transfection, cells were infected with virus for 3 h in serum free conditions, media replaced with fresh complete DMEM and maintained until harvest. For treatments with 10μM Baricitinib (Sigma-Aldrich, Cat #SBR00076), 50μM Mdivi-1(MedChemExpress, CAS 338967-87-6), 5μM Mitochondrial Division Inhibitor (MIDI) (MedChemExpress,CAS 348145-43-7), 50μM CDK1/5i (Sigma-Aldrich, CAS 40254-90-8), 2mM 2-DG (Merck, CAS 154-17-6), 2mM DL-G (Merck, CAS 56-82-6) and 25μM UK5099 (MedChemExpress, CAS 56396-35-1), the indicated inhibitors were added to complete media at 3 h post-infection and maintained for the subsequent 21 h. Cells and culture supernatants were then collected for further analysis. For pyruvate-treatment experiments, cells were pre-incubated for 24 h in glucose and pyruvate-free medium supplemented with the desired concentration of sodium pyruvate and then infected with the virus for 3 h in glucose-free, serum-free medium. After infection, the inoculum was replaced with glucose-free medium containing the corresponding pyruvate concentration, and the cells were incubated until harvesting for downstream analyses. Cells were harvested at 24 hours post infection (hpi),unless otherwise specified.

### Generation of MFN1 KO Huh-7

MFN1 KO Huh-7 cells were generated using the CRISPR-Cas9 system. Single-guide RNAs (sgRNAs) targeting exons 4 and 10 of the MFN1 gene were inserted into the PX459 vector containing WT Cas9 (Addgene, Cat #62988). Huh-7 cells were transfected with the sgRNA-containing PX459 plasmids using Lipofectamine 3000. Following transfection, puromycin selection was used to enrich transfected cells. Colonies were screened for the pure knockout lines. Knockout validation was performed by immunoblotting in multiple clones to confirm the loss of MFN1 protein. Successfully edited clones were expanded and verified as MFN1 knockout Huh-7 cells. sgRNA primers used are given in Table 3.

**TABLE 3.**
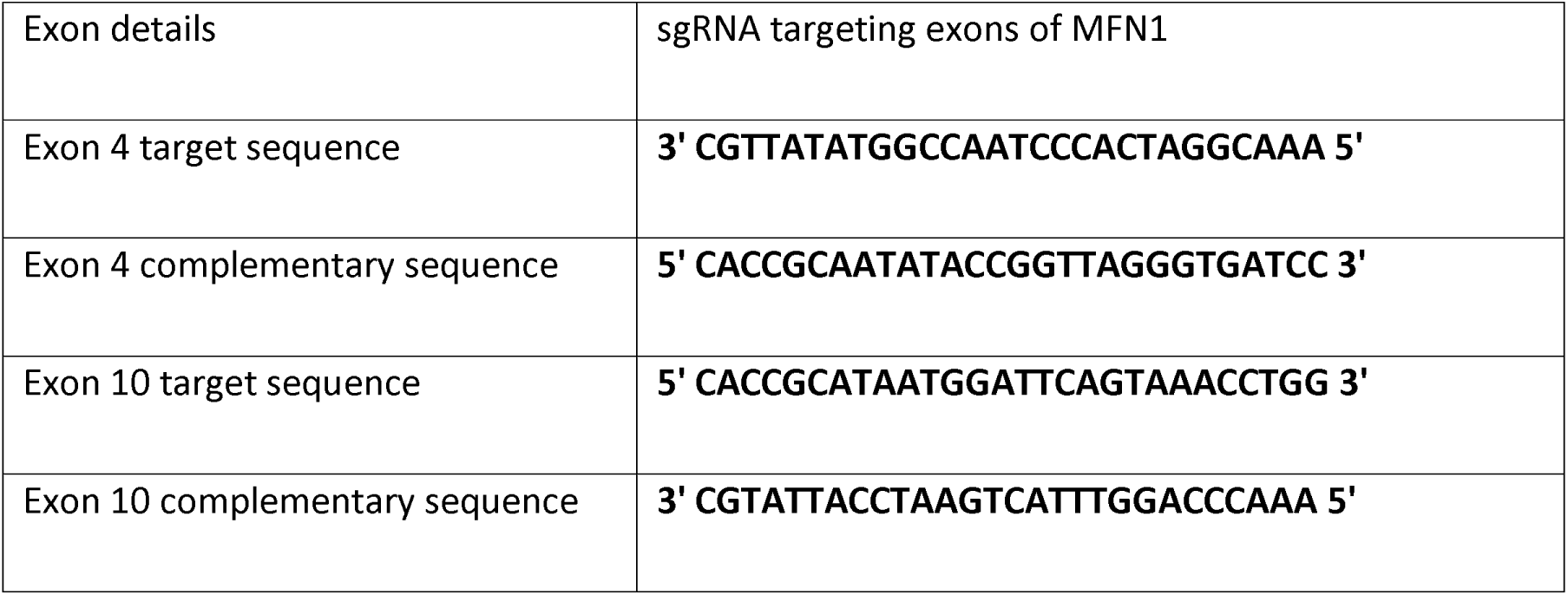
sgRNA oligos

Immu noblot ting Cell pellets were lysed

in 1× Nonidet P-40 lysis buffer containing 1% Nonidet P-40, 50 mM Tris-HCl, 150 mM NaCl (pH 7.5), EGTA, 1 mM sodium orthovanadate, 10 mM sodium pyrophosphate, 100 mM NaF, and 1 mM PMSF. Lysates were incubated on ice for 20 min with intermittent vortexing and then centrifuged at 13,000× rpm for 15 min at 4°C. Protein-containing supernatants were collected, and protein concentrations were determined using BCA reagents (Pierce, Cat #23225). Lysates were then combined with 6× denaturing loading dye, and proteins were separated by SDS-PAGE and transferred onto PVDF membranes.

Membranes were blocked with 5% BSA in 1× TBST and incubated overnight at 4°C with primary antibodies against the proteins of interest diluted in blocking buffer. After washing with 1×TBST, membranes were incubated with HRP-conjugated secondary antibodies. Immunoreactive bands were visualized using Clarity Western ECL substrate (Bio-Rad, Cat #1705061) or Clarity Max Western ECL substrate (Bio-Rad, Cat #1705062) on a Bio-Rad ChemiDoc MP system. Individual protein band intensities were quantified by densitometric analysis and normalized to the corresponding GAPDH or β- Tubulin loading control. Relative fold changes in protein expression were subsequently calculated by comparing the normalized values between experimental and control conditions and presented graphically.

Microscopy experiments MitoTracker Red staining

A total of 50,000 cells were seeded onto coverslips in 6-well plates and subjected to the indicated experimental conditions. Cells were washed twice with serum-free medium (SFM) and incubated with 100 nM MitoTracker Red CMXRos dye (CST, Cat #9082) in SFM for 20–30 min at 37°C in the dark.

Following staining, the cells were washed twice with SFM, fixed with 4% formaldehyde for 15 min at 37°C, washed with PBS, and stored at 4°C until immunostaining.

### Immunofluorescence

Fixed coverslips were rinsed twice with PBS and permeabilized for 10 min in PBS containing 0.05% Tween-20 and 0.25% Triton X-100. After three washes with PBST, samples were blocked in PBST supplemented with 1% BSA and glycine. Coverslips were incubated overnight at 4°C with primary antibodies diluted in 1% BSA/PBST. Following washing, fluorophore-conjugated secondary antibodies were added and incubated for 1 hour. Next, the coverslips were washed three additional times with PBST, and samples were mounted using DAPI-containing antifade mounting medium (VectaShield, Cat #H-1200-10) for further analysis by microscopy.

### Imaging and analysis

Confocal microscopy was performed using an Olympus FV3000 and Leica SP8 microscopes (60× oil immersion objective for Olympus FV3000, 63× oil immersion objective for Leica SP8, 2.5 × optical zoom). Mitochondrial mean branch length analysis was measured using MiNA (Mitochondrial Network Analysis) plugin inspired macros in ImageJ. Co-localization (e.g. pDRP1(S616)-mitochondria) was quantified with ImageJ EzColocalization plugin.

### Seahorse extracellular flux analysis

Glycolytic functions were assessed using a Seahorse XF Extracellular Flux Analyzer (Agilent Technologies). For the Glycolysis Stress Test, WT and MFN1 KO cells, with or without Mdivi-1 treatment, were incubated in glucose-free assay medium followed by sequential addition of 10mM glucose, 1μM oligomycin, and 50mM 2-DG according to manufacturer’s instructions. Extracellular acidification rate (ECAR) was measured and normalized to total cellular protein content (µg). Glycolysis was calculated as the difference between the maximum ECAR following glucose addition and the baseline ECAR before glucose injection. Glycolytic capacity was determined by subtracting baseline ECAR from the maximum ECAR following oligomycin addition, whereas glycolytic reserve was calculated as the difference between glycolytic capacity and glycolysis. The individual glycolytic parameters were calculated from the ECAR profiles and plotted separately for each experimental condition.

### MTT assay

Cell viability was assessed using the MTT assay. Huh7 cells were seeded in 96-well plates and treated with different concentrations of Mdivi-1, MIDI, CDK1/5i, or UK5099. Following treatment, cells were incubated with MTT reagent(500ug/well) (Sigma-Aldrich, CAS 298-93-1) and incubated for 3 hours. The resulting formazan crystals were solubilized using dimethyl sulfoxide (DMSO) for 20 mins with continuous shaking. Absorbance was measured at 570nm and reference at 650nm using a microplate reader, and cell viability was expressed as a percentage relative to untreated controls, which were normalized to 100%.

### Glycolysis assay

Glycolytic activity was assessed using the Glycolysis Assay Kit (Sigma-Aldrich, MAK439) according to the manufacturer’s instructions. Cells were cultured in low-serum (≤1% FBS) with 2.5g/L glucose and treated with Mdivi-1 or vehicle control. Following treatment, culture supernatants were collected and extracellular L-lactate, used as a measure of glycolytic activity, was quantified.

### Succinate assay

Intracellular succinate levels were measured using the Succinate Assay Kit (Sigma-Aldrich, MAK335) according to the manufacturer’s instructions. Cells treated with Mdivi-1 or vehicle control were harvested, washed with PBS, and lysed in ultrapure water. Samples were incubated with the assay reagent mixture for 30 min at room temperature, and fluorescence was measured at 530 nm excitation and 585 nm emission. Succinate concentrations were calculated from a standard curve and normalized to cell number.

### Mice experiments

All mice were bred and maintained under specific pathogen-free conditions at the CCMB Animal Facility. B6129SF2/J mice, 3-4 weeks old mice of either sex, were randomly assigned to vehicle-treated or Mdivi- 1-treated JEV-infected groups. The JEV S3 strain, kindly provided by Dr. Arup Banerjee’s group^55^, was further propagated for 2 passages in B6129SF2/J pups, and the resulting viral inoculum was titrated prior to use. Mice were infected intraperitoneally with 1 × 10^7 PFU of JEV S3. From 1-day post-infection (dpi), mice received daily intraperitoneal injections of either 50mg/kg body weight Mdivi-1 or vehicle DMSO diluted in olive oil until 5 dpi. Body weight and clinical symptoms were monitored in both groups for up to 15 dpi as previously described^56^. All animal experiments were approved by the Institutional Animal Ethics Committee of the Centre for Cellular and Molecular Biology and were conducted in accordance with institutional guidelines.

### DRP1 protein interaction network analysis

Human DRP1 (DNM1L) protein interaction data were retrieved from BioGRID (v5.0.283) on 3 November 2026. After excluding cross-species, genetic, RNA-associated, and self-interactions, redundant records were consolidated, yielding 331 interacting protein partners. Of these, 42 proteins associated with metabolism, mitochondrial dynamics, and signaling were manually selected and classified into six functional groups based on UniProt annotations. The interaction network was visualized using Python (Matplotlib), with nodes representing proteins and edges indicating reported associations. Filled and open nodes distinguish low-throughput-supported and high-throughput-only interactions, respectively, while dashed edges indicate proximity-based or co-fractionation evidence. Functional grouping was descriptive, without enrichment analysis. The figure was exported as 600dpi PNG.

## Statistical analysis

Unless otherwise specified, each experiment included at least three independent replicates. Data are presented graphically as the mean ± SEM. Statistical significance was evaluated using a two-tailed, unpaired Student’s t-test, with P-values ≤ 0.05, P ≤ 0.01, and P ≤ 0.001 denoted by *, **, and ***, respectively.

## Biosafety statement

All infection protocols in this study were approved by the Institutional Biosafety Committee of CSIR-CCMB.

## ACKNOWLEDGMENTS

We thank Dr. Joman Joseph for sharing the pEGFP-DRP1 WT plasmid. We also thank Anamta for pcDNA4/TO-MFN1 K88T cloning. We extend our thanks to Suman Bandari and Dr. Venkata Mahesh Nitla for their help with confocal imaging and analysis, and N. Sai Ram, B. Jyothi Lakshmi and Dr. M. Jerald Mahesh Kumar from CCMB Animal Facility. Special appreciation goes to Mohan Singh Moodu and Amit Kumar for their assistance with experimental logistics.

## AUTHOR CONTRIBUTIONS

P.S.P and K.H.H conceived and designed this study. P.S.P, V.S., and K.H.H. designed the experiments. P.S.P and J.S.D generated MFN1 KO cells. P.S.P, J.S.D, A.C, M.B and J.V performed kinetics experiments, inhibitor treatments, KO and overexpression studies. P.S.P and V.S performed confocal imaging and quantifications. P.S.P and P.P.S conducted metabolism-related experiments. D.T performed cell viability assays with inhibitors and conducted dual inhibition studies. V.S performed DRP1 interaction network analysis from BioGRID. P.P generated DRP1 shRNA constructs. P.S.P and K.H.H wrote the manuscript.

## Declaration of generative AI and AI-assisted technologies in the writing process

During the preparation of this work, the authors used ChatGPT to improve language and readability. After using this tool/service, the authors reviewed and edited the content as needed and take full responsibility for the content of the publication.

