## Supplemental figures 1-7 for "DRP1 Orchestrates RNA Virus Permissiveness Through Metabolic Reprogramming Independent of Mitochondrial Dynamics"

**Figure S1**

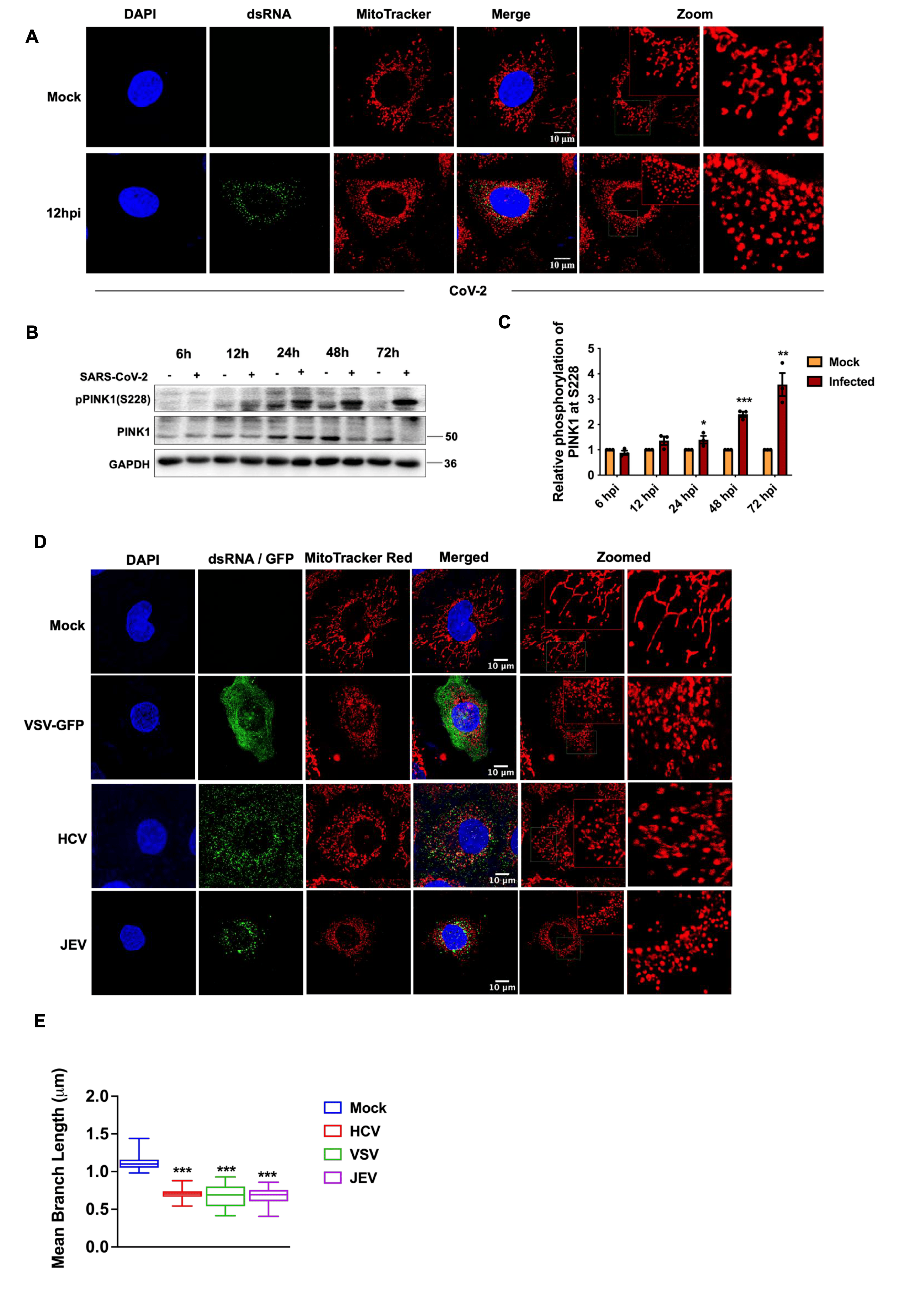

**Figure S1. Mitochondrial fission is induced during SARS-CoV-2 infection and represents a recurring feature of RNA-virus infections.**

**(A)** Confocal microscopy images showing mitochondrial fragmentation in SARS-CoV-2-infected Huh7 cells at 12 hours post-infection (hpi). Mitochondria were labeled with MitoTracker Red (red). dsRNA, stained in green, confirms viral replication.

**(B)** Immunoblots showing phosphorylated PINK1 (pPINK1-S228) and total PINK1 levels in Huh7 cells at the indicated time points following SARS-CoV-2 infection.

**(C)** Densitometric quantification of relative PINK1 phosphorylation shown in (B).

**(D)** Confocal microscopy images showing mitochondrial fragmentation in Huh7 cells infected with VSV-GFP, HCV, or JEV at 24 hpi.

**(E)** Quantification of mitochondrial mean branch length in cells shown in (D).

Scale bars, 10 µm. *n* = 50 cells unless otherwise indicated.

**Figure S2**

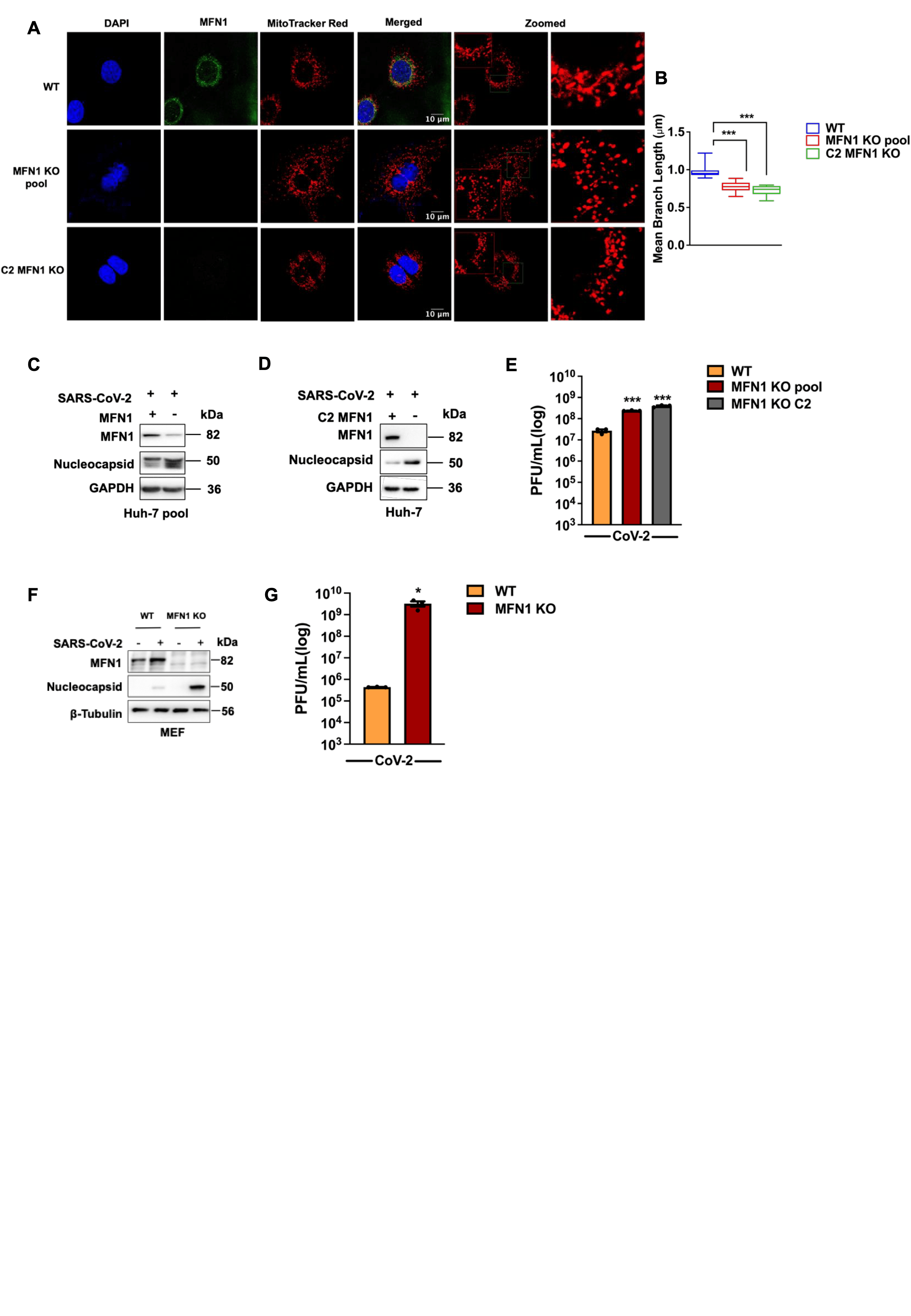

**Figure S2. Mitochondrial fission associated with MFN1 depletion enhances viral permissiveness.**

**(A)** Confocal microscopy images showing mitochondrial fragmentation in the CRISPR-Cas9-generated MFN1 KO Huh7 pooled population and an independently isolated MFN1 KO clone (C2). Mitochondria stained with MitoTracker Red(red) and MFN1 stained (green).

**(B)** Quantification of mean mitochondrial branch length in cells shown in (A).

**(C)** Representative immunoblots showing MFN1 and SARS-CoV-2 nucleocapsid (N) protein levels in wild-type (WT) Huh7 cells and the MFN1 KO pooled population following SARS-CoV-2 infection for 24 h.

**(D)** Immunoblots showing MFN1 and SARS-CoV-2 nucleocapsid (N) protein levels in WT Huh7 cells and MFN1 KO clone C2 following SARS-CoV-2 infection for 24 h.

**(E)** Infectious viral titers measured in culture supernatants from the MFN1 KO pooled population and clone C2 experiments shown in (C) and (D), respectively.

**(F)** Immunoblots showing MFN1 and SARS-CoV-2 nucleocapsid (N) protein levels in wild-type (WT) and MFN1 KO mouse embryonic fibroblasts (MEFs) following SARS-CoV-2 infection for 24 h.

**(G)** Infectious viral titers measured in culture supernatants from (F)

Scale bars, 10 µm. *n* = 50 cells unless otherwise indicated.

**Figure S3**

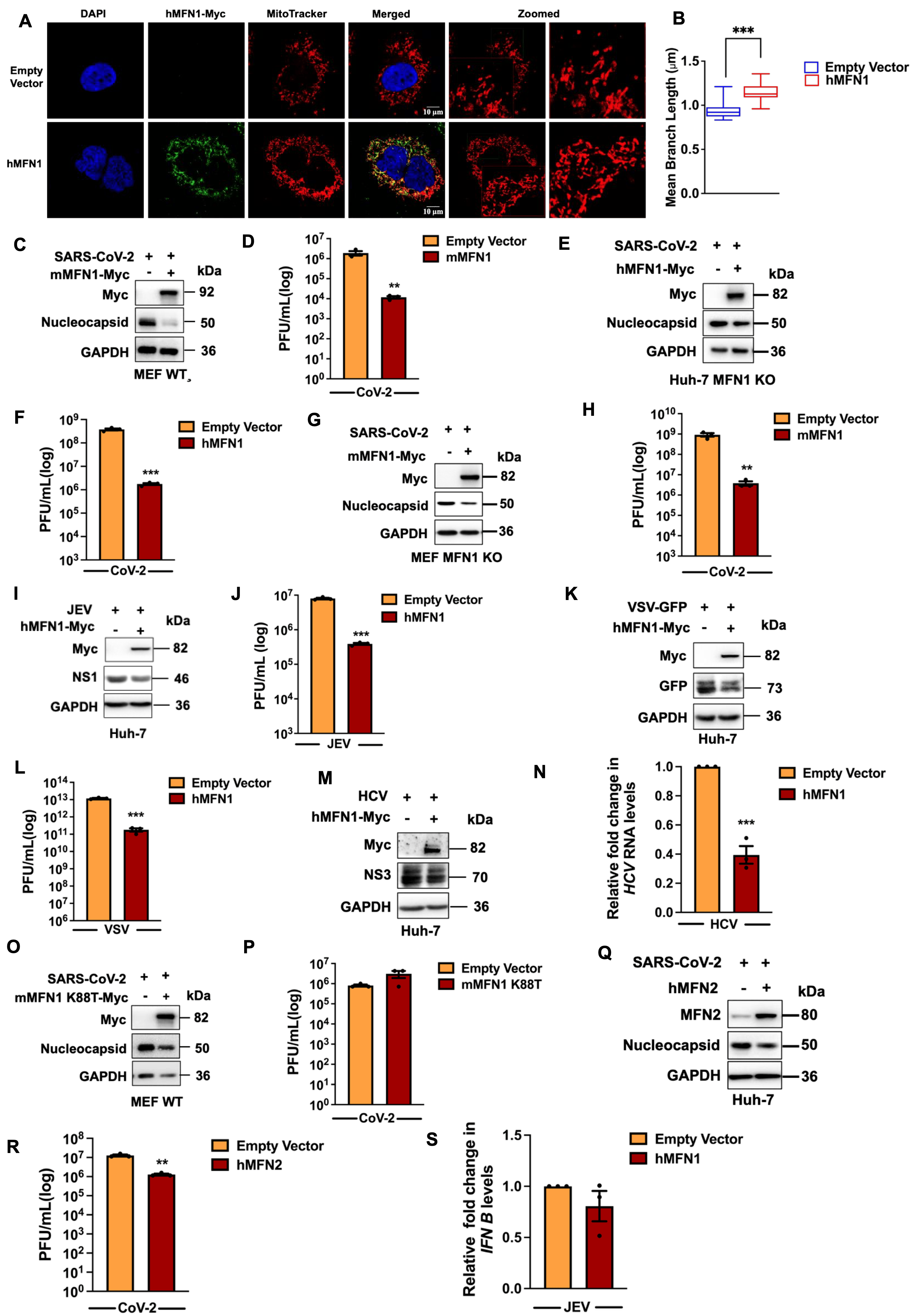

**Figure S3. Mitochondrial fusion establishes an antiviral state in host cells independently of type I interferon signaling.**

**(A)** Confocal microscopy images showing mitochondrial elongation in Huh7 cells overexpressing Myc-tagged human MFN1 (hMFN1-Myc), compared with empty vector controls. Mitochondria were labelled with MitoTracker Red (red) and Myc-tagged hMFN1 stained in (green)

**(B)** Quantification of mean mitochondrial branch length in cells shown in (A).

**(C)** Immunoblots showing Myc-tagged mouse MFN1 (mMFN1-Myc) expression and SARS-CoV-2 nucleocapsid (N) protein levels in MEF cells overexpressing mMFN1-Myc followed by SARS-CoV-2 infection.

**(D)** Infectious viral titers measured in culture supernatants from (C).

**(E)** Immunoblots showing Myc-tagged human MFN1(hMFN1-Myc) expression and SARS-CoV-2 nucleocapsid (N) protein levels in SARS-CoV-2 infected MFN1 KO Huh7 cells following re-expression of hMFN1-Myc.

**(F)** Infectious viral titers measured in culture supernatants from (E).

**(G)** Immunoblots showing Myc-tagged mouse MFN1(mMFN1-Myc) expression and SARS-CoV-2 nucleocapsid (N) protein levels in SARS-CoV-2 infected MFN1 KO MEFs following re-expression of mMFN1-Myc.

**(H)** Infectious viral titers measured in culture supernatants from (G).

**(I)** Immunoblots showing Myc-tagged human MFN1(hMFN1-Myc) expression and JEV NS1 protein levels in hMFN1-Myc-overexpressing Huh7 cells following JEV infection.

**(J)** Corresponding infectious viral titers measured in culture supernatants from (I).

**(K)** Immunoblots showing Myc-tagged human MFN1(hMFN1-Myc) expression and VSV-GFP levels in Huh7 cells overexpressing hMFN1-Myc following VSV infection.

**(L)** Infectious viral titers measured in culture supernatants from (K).

**(M)** Immunoblots showing Myc-tagged human MFN1(hMFN1-Myc) expression and HCV NS3 protein levels in hMFN1-Myc-overexpressing Huh7 cells following HCV infection.

**(N)** Quantification of HCV RNA levels by RT-qPCR in culture supernatants from (M).

**(O)** Immunoblots showing Myc-tagged GTPase-deficient mouse MFN1 (mMFN1-K88T-Myc) expression and SARS-CoV-2 nucleocapsid (N) protein levels in MEFs expressing the mMFN1-K88T mutant following SARS-CoV-2 infection.

**(P)** Infectious viral titers measured in culture supernatants from (O).

**(Q)** Immunoblots showing human MFN2 (hMFN2) and SARS-CoV-2 nucleocapsid (N) protein levels in hMFN2-overexpressing Huh7 cells.

**(R)** Corresponding infectious viral titers measured in culture supernatants from (Q).

**(S)** RT-qPCR analysis of *IFNB1* mRNA levels in JEV-infected Huh7 cells overexpressing hMFN1-Myc compared with empty vector controls.

Scale bars, 10 µm. *n* = 50 cells unless otherwise indicated.

**Figure S4**

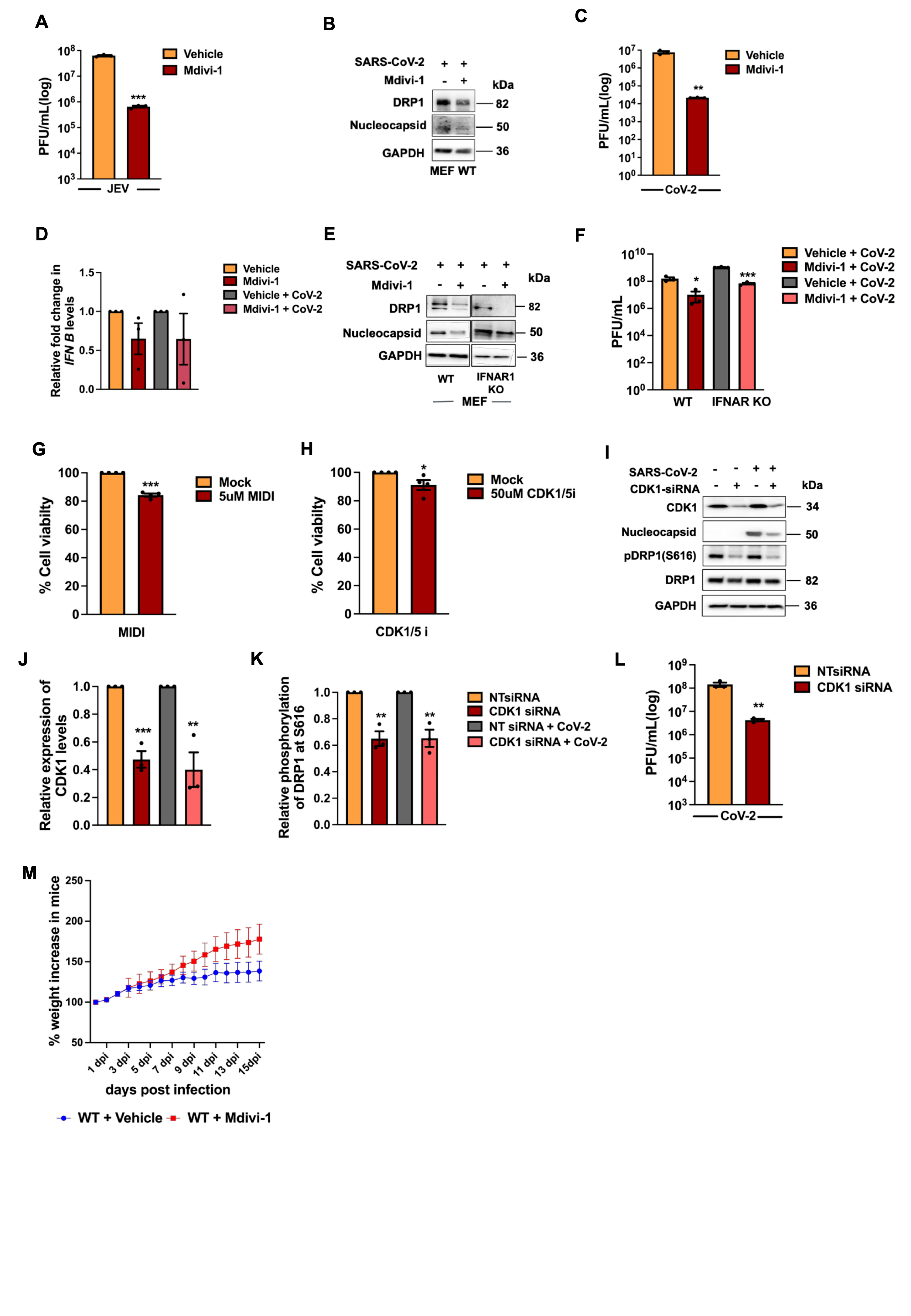

**Figure S4. Inhibition of DRP1 function establishes a potent antiviral state.**

**(A)** Infectious viral titers measured in culture supernatants from JEV-infected Huh7 cells treated with Mdivi-1 (50 µM).

**(B)** Immunoblots showing DRP1 and SARS-CoV-2 nucleocapsid (N) protein levels in mouse embryonic fibroblasts (MEFs) following SARS-CoV-2 infection and Mdivi-1 treatment (50 µM).

**(C)** Infectious viral titers measured in culture supernatants from (B).

**(D)** RT-qPCR analysis of *IFNB1* mRNA levels in uninfected and SARS-CoV-2-infected Huh7 cells treated with Mdivi-1, compared with their respective vehicle controls.

**(E)** Immunoblots showing DRP1 and SARS-CoV-2 nucleocapsid (N) protein levels in wild-type (WT) and IFNAR1-knockout (KO) primary MEFs following SARS-CoV-2 infection and Mdivi-1 treatment. Immunoblot images from WT and IFNAR1 KO cells were cropped and assembled into a single panel for presentation.

**(F)** Infectious viral titers measured in culture supernatants from (E).

**(G)** Percentage cell viability in untreated and 5µM MIDI treated Huh-7 cells, as assessed by an MTT assay.

**(H)** Percentage cell viability in untreated and 50µM CDK1/5i treated Huh-7 cells, as assessed by an MTT assay.

**(I)** Immunoblots showing CDK1, phosphorylated DRP1 (pDRP1-S616), total DRP1, and SARS-CoV-2 nucleocapsid (N) protein levels in Huh7 cells following CDK1-specific siRNA transfection and SARS-CoV-2 infection.

**(J)** Densitometric quantification of CDK1 protein levels shown in (I).

**(K)** Densitometric quantification of relative DRP1 phosphorylation at S616 shown in (I).

**(L)** Infectious viral titers measured in culture supernatants from (I).

**(M)** Percentage change in body weight of JEV-infected mice following Mdivi-1 treatment compared with untreated infected controls, monitored through 15 days post-infection (dpi).

**Figure S5**

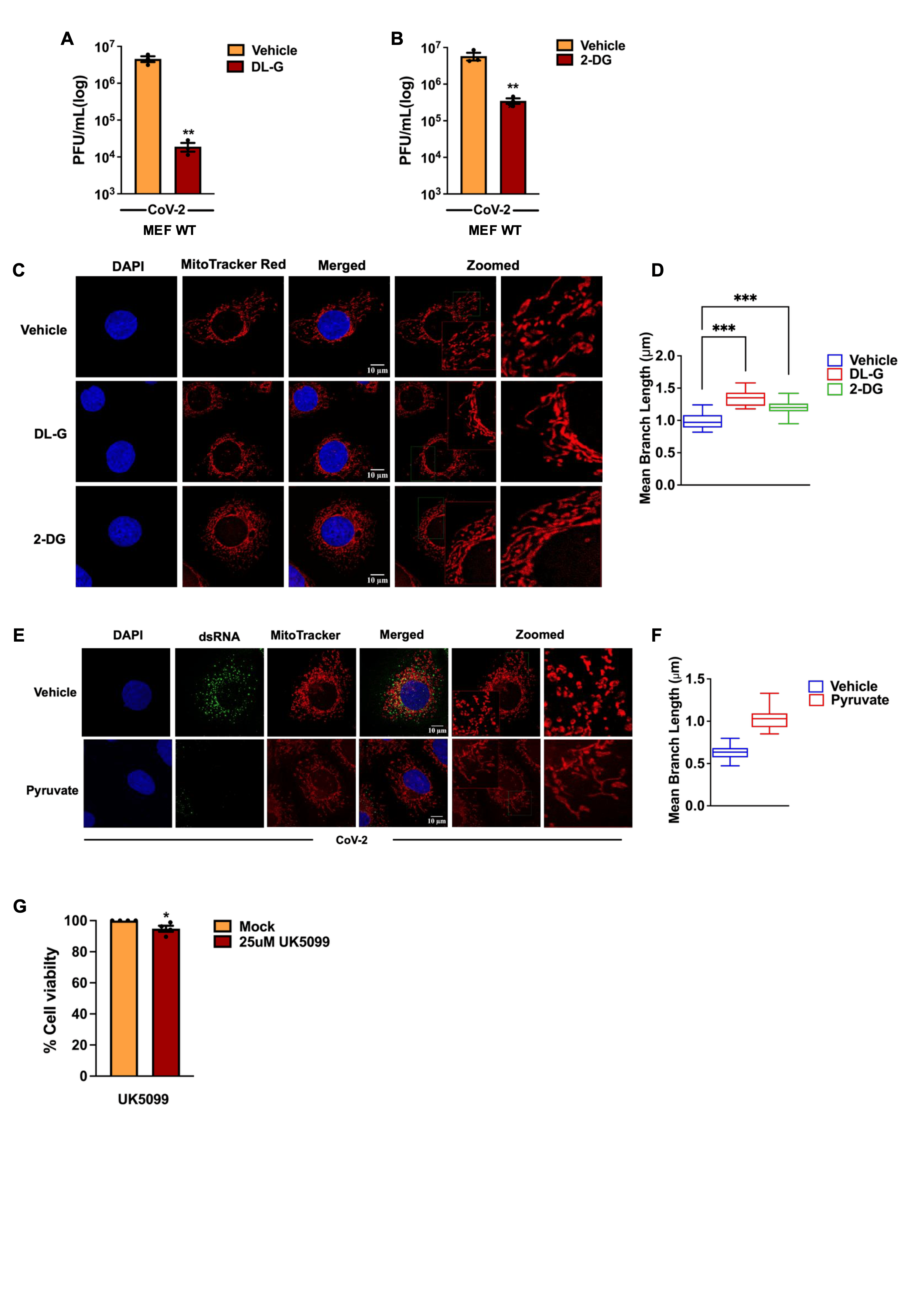

**Figure S5. A major metabolic reprogramming driven by DRP1 inhibition and mitochondrial elongation supports the antiviral state.**

**(A)** Infectious viral titers measured in culture supernatants from SARS-CoV-2-infected mouse embryonic fibroblasts (MEFs) treated with DL-glyceraldehyde (DL-G), compared with vehicle controls.

**(B)** Infectious viral titers measured in culture supernatants from SARS-CoV-2-infected MEFs treated with 2-deoxyglucose (2-DG), compared with vehicle controls.

**(C)** Confocal microscopy images showing mitochondrial morphology in uninfected Huh7 cells treated with DL-G or 2-DG, compared with their respective vehicle controls.

**(D)** Quantification of mean mitochondrial branch length in cells shown in (C).

**(E)** Representative confocal microscopy images showing mitochondrial morphology in SARS-CoV-2 infected Huh7 cells supplemented with sodium pyruvate (440 mg/L), compared with vehicle controls.

**(F)** Quantification of mean mitochondrial branch length in cells shown in (E).

**(G)** Percentage cell viability in untreated and 25µM UK5099 treated Huh-7 cells, as assessed by an MTT assay.

Scale bars, 10 µm. *n* = 50 cells unless otherwise indicated.

**Figure S6**

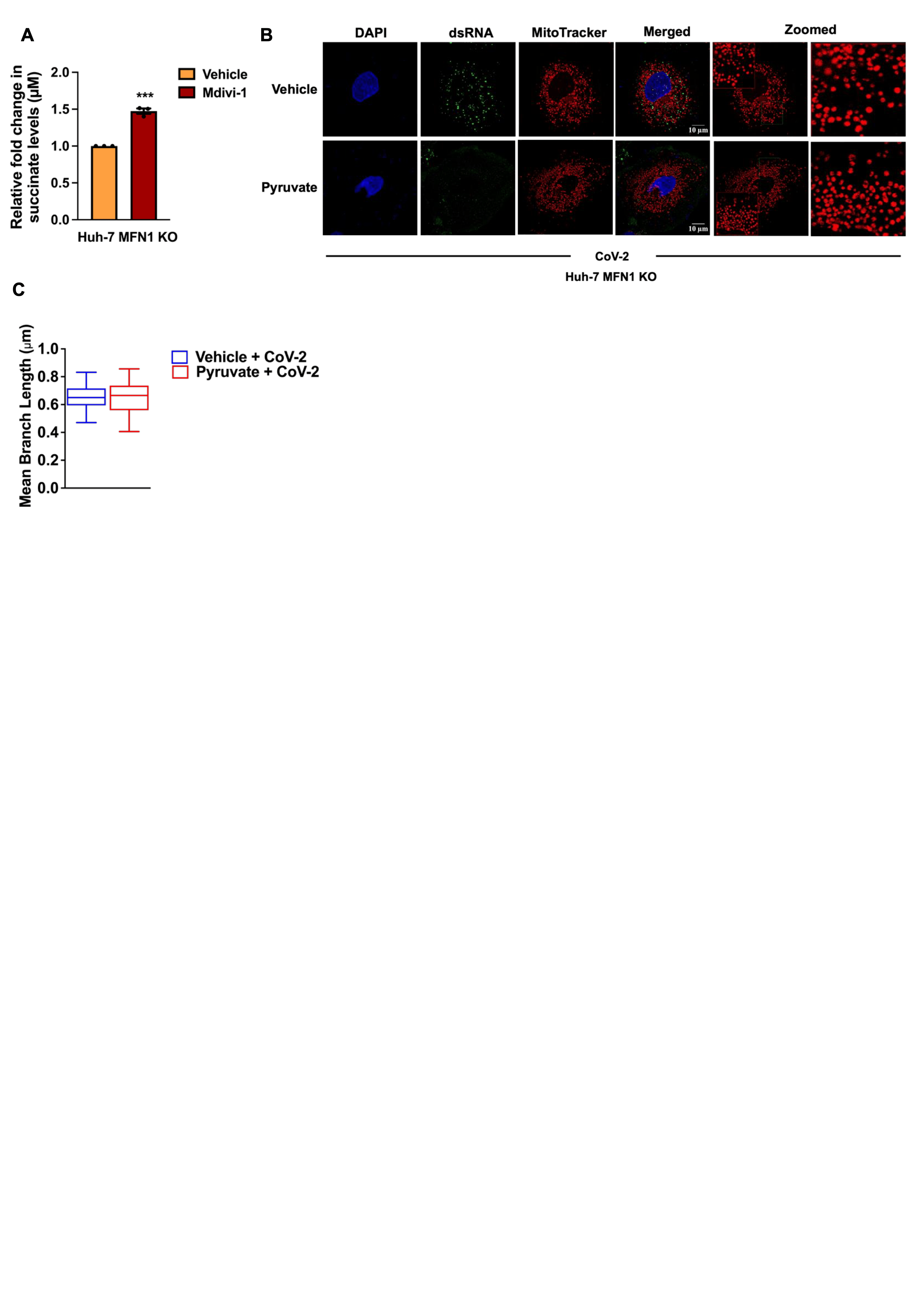

**Figure S6. MFN1 and mitochondrial fusion are dispensable for DRP1 mediated viral permissiveness.**

**(A)** Intracellular succinate levels measured in MFN1 KO Huh7 cells treated with Mdivi-1, compared with vehicle controls.

**(B)** Confocal microscopy images showing mitochondrial morphology in SARS-CoV-2-infected MFN1 KO Huh7 cells supplemented with sodium pyruvate (440 mg/L), compared with vehicle controls.

**(C)** Quantification of mean mitochondrial branch length in cells shown in (B).

Scale bars, 10 µm. *n* = 50 cells unless otherwise indicated.

**Figure S7**

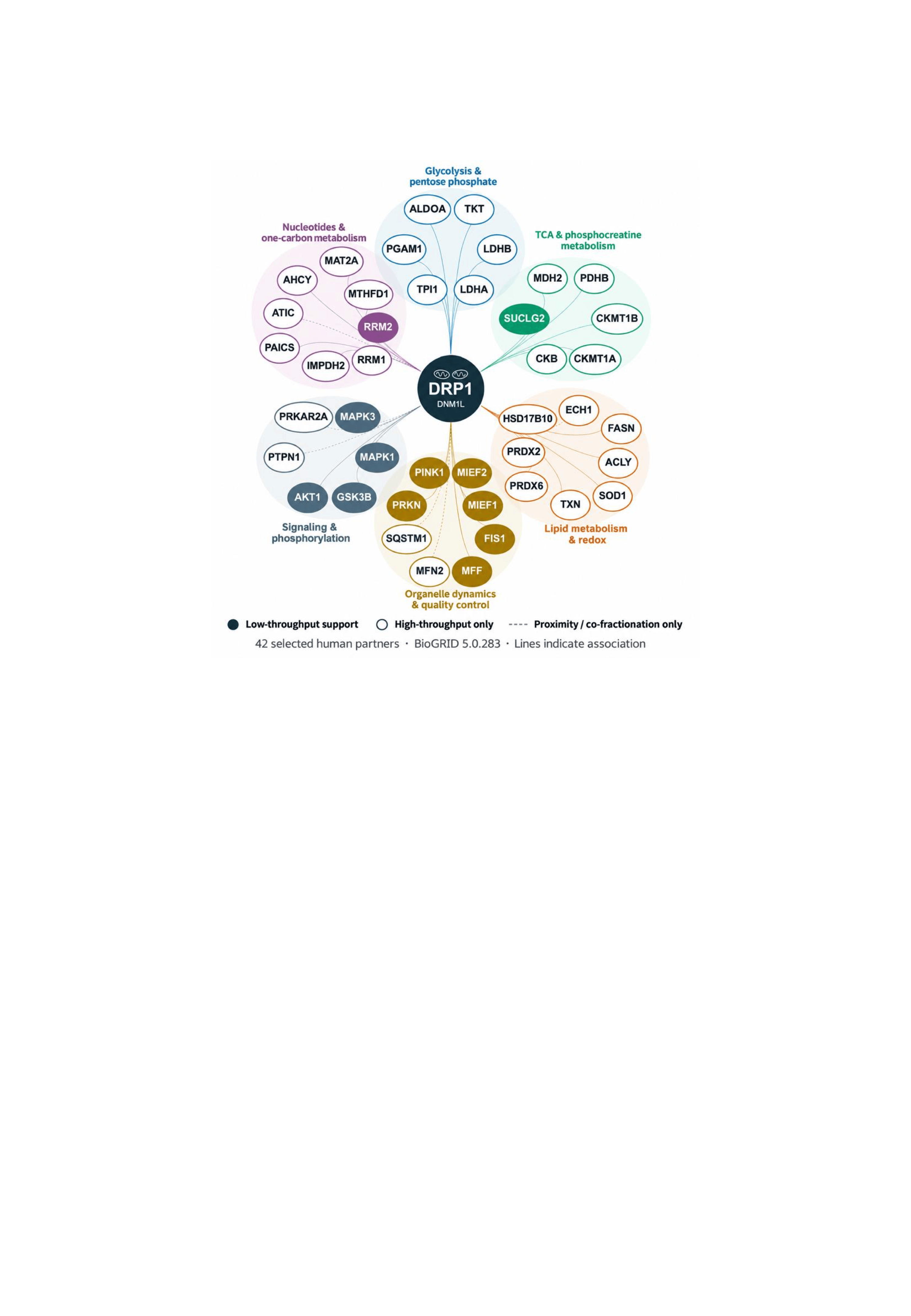

**Figure S7. DRP1-mediated viral permissiveness encompasses broader mechanisms beyond glycolytic metabolism**

Protein association network of human DRP1 (DNM1L) showing 42 selected proteins identified using BioGRID (version 5.0.283), grouped according to their roles in metabolism, mitochondrial dynamics, quality control, and signaling. Filled nodes indicate low-throughput experimental support, while unfilled nodes represent high-throughput-only evidence. Dashed lines indicate associations supported only by proximity or co-fractionation evidence. These associations do not necessarily represent direct physical interactions or functional regulation by DRP1.
